# The Arabidopsis E3 ubiquitin ligase PUB4 regulates BIK1 homeostasis and is targeted by a bacterial type-III effector

**DOI:** 10.1101/2020.10.25.354514

**Authors:** Maria Derkacheva, Gang Yu, Jose S. Rufian, Shushu Jiang, Paul Derbyshire, Rafael J. L. Morcillo, Lena Stransfeld, Yali Wei, Frank L.H. Menke, Cyril Zipfel, Alberto P. Macho

## Abstract

Plant immunity is tightly controlled by a complex and dynamic regulatory network, which ensures optimal activation upon detection of potential pathogens. Accordingly, each component of this network is a potential target for manipulation by pathogens. Here, we report that RipAC, a type III-secreted effector from the bacterial pathogen *Ralstonia solanacearum*, targets the plant E3 ubiquitin ligase PUB4 to inhibit pattern-triggered immunity (PTI). PUB4 plays a positive role in PTI by regulating the homeostasis of the central immune kinase BIK1. Before PAMP perception, PUB4 promotes the degradation of non-activated BIK1, while, after PAMP perception, PUB4 contributes to the accumulation of activated BIK1. RipAC leads to BIK1 degradation, which correlates with its PTI-inhibitory activity. RipAC causes a reduction in pathogen-associated molecular pattern (PAMP)-induced PUB4 accumulation and phosphorylation. Our results shed light on the role played by PUB4 in immune regulation, and illustrate an indirect targeting of the immune signalling hub BIK1 by a bacterial effector.

## Introduction

Plants constantly face pathogens, and the success of their defence depends on their ability to sense an attack and provide a fast response. The first layer of plant immunity is based on the recognition of pathogen-associated molecular patterns (PAMPs) or damaged-associated molecular patterns (DAMPs) via plasma membrane-localized pattern recognition receptors (PRRs). Perception of PAMPs or DAMPs by their corresponding PRRs leads to pattern-triggered immunity (PTI), which restricts the multiplication of most potential pathogens (Couto and Zipfel, 2016). The *Arabidopsis thaliana* (hereafter Arabidopsis) leucine-rich repeat receptor kinases (LRR-RKs) FLS2 and EFR are amongst the best-characterized plant PRRs. They recognize the bacterial PAMPs flagellin (or its derived peptide flg22) and elongation factor Tu (or its derived peptide elf18), respectively (Gómez-Gómez and Boller, 2000; Zipfel et al., 2006). The plant elicitor peptide 1 (AtPep1), which is secreted by plant cells to amplify the immune response, is perceived by the LRR-RKs PEPR1/PEPR2 (Krol et al., 2010; Tang and Zhou, 2015; Yamaguchi et al., 2010; Yamaguchi et al., 2006). Ligand-binding triggers the instantaneous association between FLS2/EFR/PEPR1 and the LRR-RK BAK1 (also known as SERK3) and related SERKs, which act as co-receptors, leading to phosphorylation events and activation of downstream immune signalling (Chinchilla et al., 2007; Heese et al., 2007; Perraki et al., 2018; Postel et al., 2010; Roux et al., 2011; Schulze et al., 2010; Schwessinger et al., 2011; Sun et al., 2013). Chitin, a component of the fungal cell wall, is perceived by the LysM-RK LYK5, which forms a complex with the co-receptor CERK1 (another LysM-RK) upon ligand-binding, leading to trans-phosphorylation and initiation of immune signalling (Cao et al., 2014; Erwig et al., 2017; Liu et al., 2018; Liu et al., 2012; Miya et al., 2007; Petutschnig et al., 2010; Suzuki et al., 2016; Suzuki et al., 2018; Suzuki et al., 2019; Wan et al., 2008).

The activation of PRR complexes leads to the activation of downstream receptor-like cytoplasmic kinases (RLCKs) (Liang and Zhou, 2018). The RLCK BIK1 is a direct substrate of FLS2/EFR/PEPR1/CERK1 complexes and is thus a convergent point in PTI signalling triggered by several elicitors (Laluk et al., 2011; Liu et al., 2013; Lu et al., 2010; Zhang et al., 2010). BIK1 plays roles in plant immunity to bacterial and fungal pathogens (Lu et al., 2010; Veronese et al., 2006; Zhang et al., 2010), and is required for several PTI responses, such as production of reactive oxygen species (ROS), calcium influx, callose deposition, and stomata closure (Kadota et al., 2014; Li et al., 2014; Lu et al., 2010; Monaghan et al., 2015; Ranf et al., 2014; Thor et al., 2020; Tian et al., 2019; Zhang et al., 2010).

BIK1 is a rate-limiting factor in PTI responses, and as such BIK1 protein levels are tightly regulated (Couto and Zipfel, 2016; Kadota et al., 2014; Jiang et al., 2019; Liang et al., 2016; Monaghan et al., 2014; Wang et al., 2018; Zhang et al., 2010; Zhang et al., 2018). It was suggested that two pools of BIK1 exist in a cell: ‘non-activated’ (non-phosphorylated, before PAMP treatment) and ‘activated’ (phosphorylated, after PAMP treatment) (Wang et al., 2018). Non-activated BIK1 is targeted for proteasomal degradation by the E3 ubiquitin ligases PUB25 and PUB26, which is promoted by the cytoplasmic calcium-dependent kinase CPK28; on the contrary, this degradation is inhibited by the heteromeric G protein complex formed by XLG2-AGB1-AGG1/2 (Liang et al., 2016; Monaghan et al., 2014; Wang et al., 2018). After flg22 perception, the heteromeric G protein complex dissociates from FLS2, and CPK28 phosphorylates PUB25/26 to promote degradation of the non-activated pool of BIK1, and to adjust the amplitude of immune response (Liang et al., 2016; Wang et al., 2018). Activated BIK1 dissociates from the FLS2-BAK1 complex and is protected from degradation by PUB25/26 while promoting immune signalling (Wang et al., 2018).

To achieve a successful infection, pathogens secrete effector proteins to manipulate plant cellular functions, including the targeting of immune signalling components and the manipulation of PTI (Lee et al., 2019; Macho, 2016; Macho and Zipfel, 2015). Therefore, besides being important virulence factors, effector proteins constitute useful probes to identify and characterize novel plant proteins involved in immune signalling (Toruño et al., 2016). *Ralstonia solanacearum* is a soil-borne pathogen, and causal agent of bacterial wilt disease. *R. solanacearum* can infect more than 250 plant species that belong to more than 50 families, including economically important crops such as tomato, potato, banana, pepper and eggplant (Jiang et al., 2017; Mansfield et al., 2012; Wicker et al., 2007). *R. solanacearum* enters the plants through the roots, reaches the vascular system, and proliferates in xylem vessels to colonize the whole plant (Genin, 2010; Xue et al., 2020). *R. solanacearum* relies on a type-III secretion system to deliver over 70 Type-III effector proteins (T3Es) into the host cytoplasm (Sabbagh et al., 2019). One of these T3Es, RipAC, is able to suppress effector-triggered immunity (ETI) by targeting SGT1, and is required for full virulence of *R. solanacearum* in tomato and Arabidopsis (Yu et al., 2020a).

Here, we demonstrate that RipAC suppresses diverse PAMP-induced responses. We also show that RipAC interacts with the E3 ubiquitin ligase PUB4 from tomato and Arabidopsis, and that *pub4* mutant plants show deficient immune responses to several PAMPs. Interestingly, we found that PUB4 has a dual impact on the early PTI regulator BIK1: before PAMP treatment, PUB4 promotes degradation of non-activated BIK1, but, after PAMP treatment, PUB4 is required for the accumulation of activated BIK1. RipAC overexpression in Arabidopsis leads to BIK1 degradation, suggesting that RipAC exploits PUB4 to degrade BIK1 and suppress PTI responses.

## Results

### RipAC suppresses pattern-triggered immunity

To verify whether RipAC affects PTI in Arabidopsis, we characterized PAMP-induced responses in Arabidopsis transgenic lines overexpressing RipAC (Yu et al., 2020a). Two independent *RipAC-GFP* overexpression lines showed a decreased ROS burst in response to bacterial or fungal PAMPs (flg22^Pto^, elf18^Rsol^, and chitin; Figures 1A-C). The level of ROS inhibition correlated with the level of RipAC accumulation in these lines (Figure 1D). Activation of PAMP-induced signalling leads to the activation of MAPK cascades (Yu et al., 2017). Flg22-induced MAPK activation was also decreased in *RipAC-GFP* overexpression lines (Figure 1D). In correspondence with the observed inhibition of early PTI responses, *RipAC* overexpression lines were more susceptible to *Pseudomonas syringae* pv. tomato (*Pto)* DC3000 *ΔhrcC*, a non-pathogenic mutant strain unable to secrete T3Es, and therefore inducing solely PTI, but not to wild-type *Pto* DC3000, which is able to suppress PTI (Figure 1E). Together, these results demonstrate that RipAC inhibits early PTI responses triggered by various PAMPs to facilitate pathogen infection.

**Figure 1.**
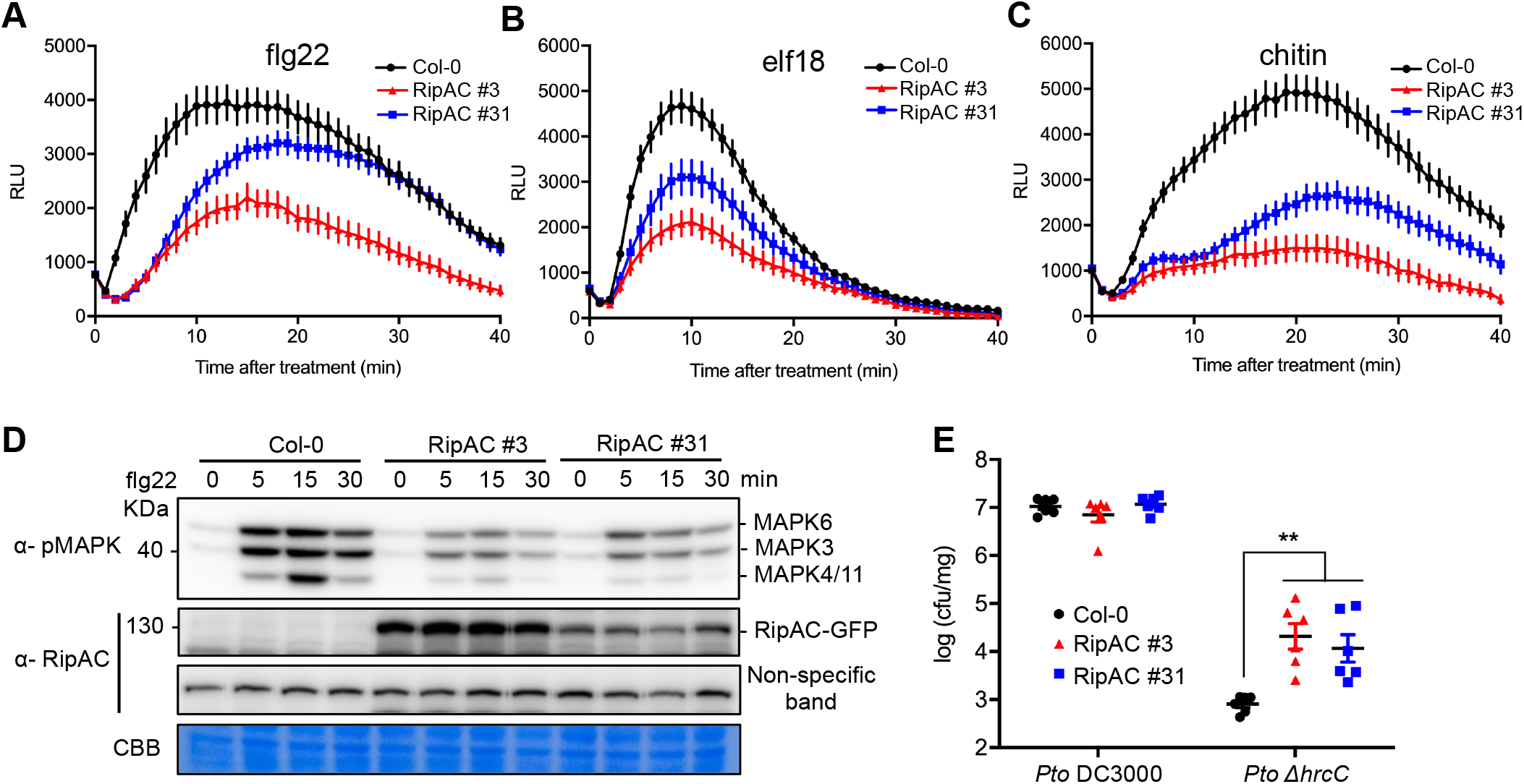
RipAC inhibits PTI in Arabidopsis. (A-C) RipAC overexpression inhibits ROS production induced by 50 nM flg22^Pto^ (A), 100 nM elf18^Rsol^ (B), and 200 mg/mL chitin (C) in Arabidopsis. ROS was measured as relative luminescence units (RLU) over time. Values are means ± SE (n=24). (D) RipAC overexpression inhibits flg22-triggered MAPK activation in Arabidopsis. 100 nM flg22^Pto^ was used to treat Arabidopsis seedlings and the samples were collected at indicated time points. Immunoblots were analysed using anti-pMAPK and anti-RipAC antibodies. Coomassie brilliant blue (CBB) staining and a non-specific band were used as loading control. Molecular weight (kDa) marker bands are indicated for reference. (E) RipAC overexpression lines display elevated susceptibility to *Pto* DC3000 Δ*hrcC*, but not to *Pto* DC3000. Arabidopsis plants were spray-inoculated with indicated *Pto* strains and bacterial titers were determined 3 days post-inoculation. Values are means ± SE (n=6). Asterisks indicate significant differences compared to Col-0 (Student’s t test, ** p<0.01). Experiments were performed 3 times with similar results.

### RipAC interacts with PUB4 from tomato and Arabidopsis

Our previous work showed that SGT1 targeting by RipAC does not underlie PTI suppression (Yu et al., 2020b), and therefore the relevant target(s) of RipAC in this context remain to be identified. As tomato (*Solanum lycopersicum*) is a major crop affected by *R. solanacearum*, we performed a yeast two-hybrid screen using RipAC as a bait against a library of cDNA from tomato roots inoculated with *R. solanacearum*. We identified several clones matching the tomato ortholog of Arabidopsis *PLANT U-BOX PROTEIN 4* (*AtPUB4*), *SlPUB4* (Figure S1A). We further confirmed the interaction between RipAC and PUB4 *in planta* using split-luciferase (Split-LUC) complementation assays in *Nicotiana benthamiana*, which showed that RipAC interacts with both SlPUB4 and AtPUB4, but not with the aquaporin AtPIP2A used as negative control (Figures 2A, B, and S1B). Co-expression of GFP-tagged SlPUB4/AtPUB4, or GFP alone, with RipAC-nLUC in *N. benthamiana*, followed by co-immunoprecipitation (CoIP), revealed RipAC association with SlPUB4 and AtPUB4 (Figure 2C). Additionally, FRET-FLIM assays in *N. benthamiana* further confirmed the direct interaction of RipAC with SlPUB4 and AtPUB4 (Figure 2D). Altogether, these results indicate that PUB4 is a novel interactor of RipAC in plant cells.

**Figure 2.**
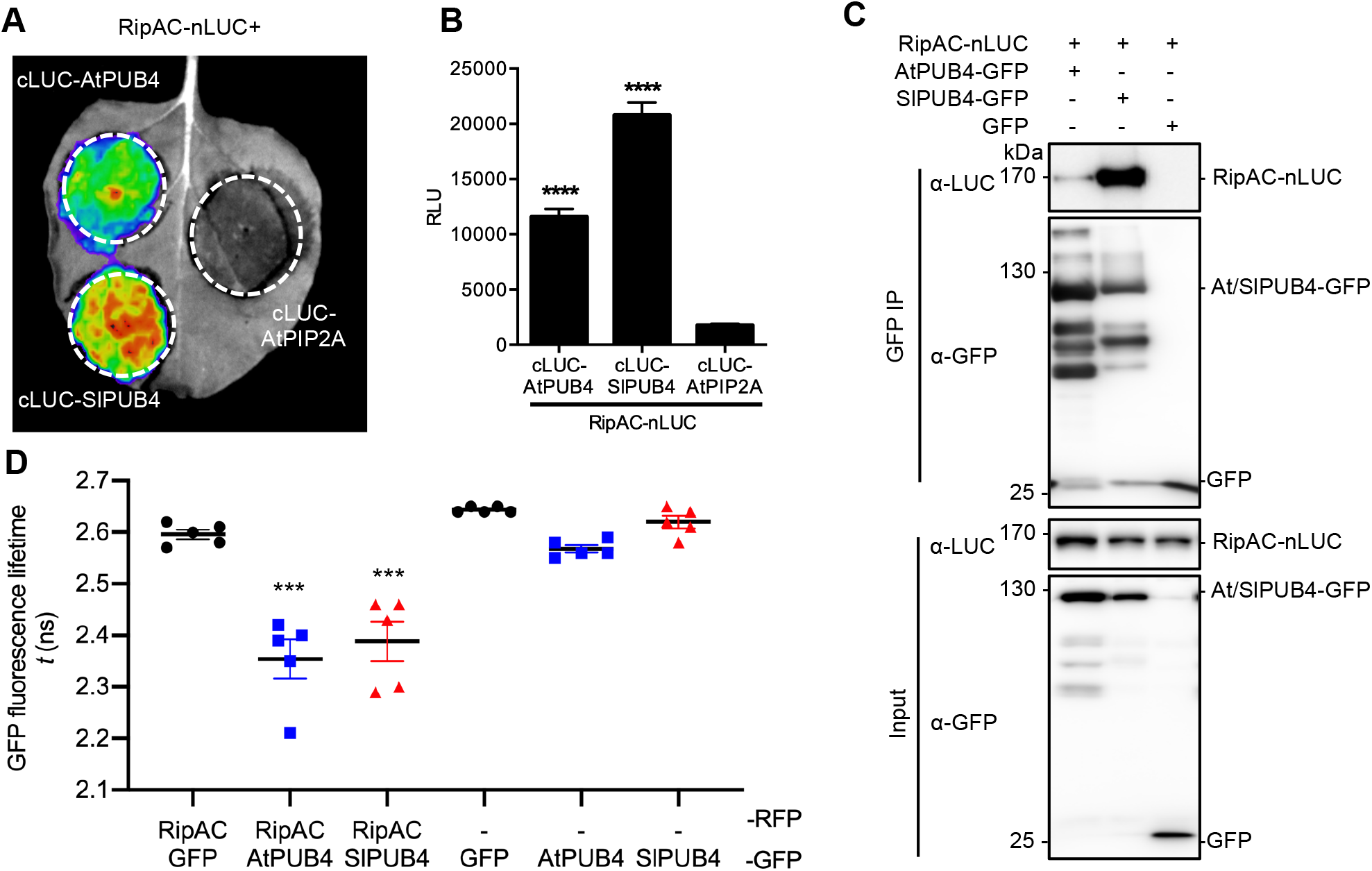
RipAC associates with PUB4 in plant cells. (A-B) Interaction between RipAC and SlPUB4/AtPUB4 was confirmed by Split-LUC assay qualitatively (A) and quantitatively (B) in *Nicotiana benthamiana*. Values are means ± SE (n=8). Asterisks indicate significant differences compared to RipAC-nLUC/cLUC-AtPIP2A negative control (Student’s t test, **** p<0.0001). RLU: relative luminescence units. (C) Co-immunoprecipitation of SlPUB4/AtPUB4-GFP and RipAC-nLUC after co-expression in *N. benthamiana*. Two days after Agrobacterium infiltration, the plant tissues were collected and then subjected to anti-GFP immunoprecipitation. Immunoblots were analyzed using anti-LUC and anti-GFP antibodies. Molecular weight (kDa) marker bands are indicated for reference. (D) Interaction between RipAC-RFP and SlPUB4/AtPUB4-GFP determined by FRET-FLIM upon transient co-expression in *N. benthamiana* leaves. GFP fluorescence lifetime of each - GFP fusion protein is shown as a negative control. Lines represent average values (n=5) and error bars represent standard error. (Student’s t-test, *** p<0.001). Experiments were performed 3 times with similar results.

### PUB4 positively regulates PTI in Arabidopsis

Arabidopsis PUB4 belongs to the family of plant U-box proteins containing Armadillo (ARM) repeats (Mudgil et al., 2004). AtPUB4 has been implicated in several processes, such as cytokinin responses, tapetum development, meristem maintenance, cell division, oxidative stress, and responses to chitin (Desaki et al., 2019; Kinoshita et al., 2015a; Kinoshita et al., 2015b; Wang et al., 2017; Woodson et al., 2015).

We sought to determine whether the targeting of PUB4 might underlie the suppression of PTI responses by RipAC. First, to understand the role of PUB4 in the regulation of PTI, we characterized PAMP/DAMP-triggered responses in *pub4* mutants. Two independent *pub4* mutant lines showed reduced ROS burst in response to flg22^Paer^, elf18^Ecol^, AtPep1, and chitin (Figures 3A-D). Similar results were obtained in *pub4* mutants using flg22^Pto^ and elf18^Rsol^ as elicitors (Figures S2A-B). These results are consistent with the recently reported reduced chitin-induced ROS production in *pub4* mutant (Desaki et al., 2019). The inhibition of ROS triggered by various PAMPs in *pub4* mutants resembled the phenotype of RipAC-overexpression lines (Figure 1). MAPK activation was not affected in *pub4* mutants (Figure 3E), suggesting that the effect of RipAC on MAPK activation is PUB4-independent. The inhibition of seedling growth triggered by flg22 and elf18 was also compromised in *pub4* mutants (Figures 3F and 3G), indicating that PUB4 is also required for late PTI responses. Moreover, *pub4* mutant lines showed compromised stomatal closure triggered by flg22 or chitin (Figures 3H and 3I). Abscisic acid (ABA)-triggered stomatal closure was comparable in Col-0 and *pub4* plants (Figure 3J), indicating that *pub4* stomatal closure is specifically impaired in response to PAMPs. Finally, we tested the role of PUB4 in antibacterial immunity by performing surface-inoculation with the non-pathogenic strain *Pto* DC3000 Δ*hrcC*. Given that *pub4* mutants showed deficient PAMP-triggered stomatal closure, we also performed inoculations with a weakly virulent *Pto* derivative unable to produce coronatine (*Pto* DC3000 COR^−^), a non-T3E virulence factor required for the suppression of stomatal defences (Melotto et al., 2006). We found that *pub4* mutants were more susceptible to both strains (Figure 3K). Together, these results demonstrate that PUB4 genetically behaves as a positive regulator of PTI.

**Figure 3.**
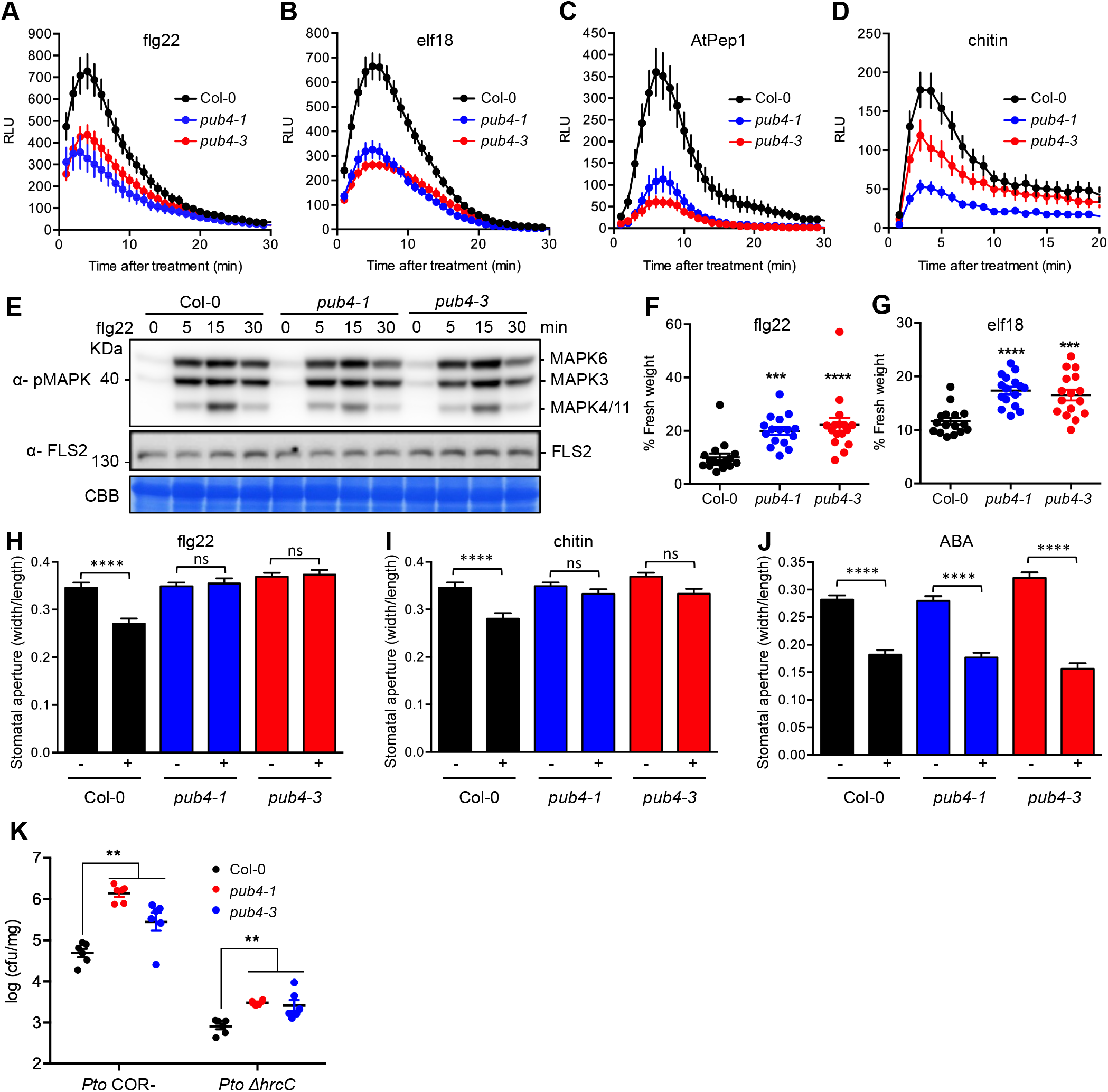
PUB4 positively regulates PTI in Arabidopsis. (A-D) ROS burst in Col-0 WT or the indicated *pub4* mutant lines induced by 100 nM flg22^Paer^ (A), 100 nM elf18^Ecol^ (B), 1 mM AtPep1 (C) and 1 mg/mL chitin (D). ROS was measured as relative luminescence units (RLU) over time. Values are means ± SE (n=16). (E) flg22-triggered MAPK activation in *pub4* T-DNA mutants. 100 nM flg22^Pto^ was used to treat Arabidopsis seedlings and the samples were collected at indicated time points for western blots. Immunoblots were analyzed using anti-pMAPK and anti-FLS2 antibodies. Coomassie brilliant blue (CBB) staining was used as loading control. Molecular weight (kDa) marker bands are indicated for reference. (F-G) PUB4 contributes to seedling growth inhibition induced by 100 nM flg22^Paer^ (A) or 100 nM elf18^Ecol^ (B). Values represent the percentage of fresh weight of PAMP-treated vs water-treated seedlings, and are means ± SE (n=16). Asterisks indicate significant differences compared to Col-0 (one-way ANOVA, Kruskal-Wallis test, Dunn’s multiple comparisons test, *** p<0.001, ****p<0.0001). (H-J) PUB4 contributes to PAMP-induced stomatal closure. Stomatal apertures were measured as width to length ratio 2 h after treatment with mock or 10 mM flg22^Paer^ (H), 1 mg/mL chitin (I) and 10 mM ABA (J). Values are mean ± SE (n>120). Asterisks indicate significant differences between samples (one-way ANOVA, Kruskal-Wallis test, Dunn’s multiple comparisons test, ****p<0.0001, ^ns^p>0.05). (K) *pub4* mutant lines display elevated susceptibility to *Pto* DC3000 COR^*−*^ and *Pto* DC3000 Δ*hrcC*. Arabidopsis plants were spray-inoculated with indicated *Pto* strains and bacterial titers were determined 3 days post-inoculation. Values are means ± SE (n=6). Asterisks indicate significant differences compared to Col-0 (Student’s t test, ** p<0.01). Experiments were performed 3 times with similar results.

### PUB4 promotes *R. solanacearum* infection in Arabidopsis and tomato

To determine whether PUB4 is required for plant basal resistance to *R. solanacearum*, we performed soil-drenching inoculation with *R. solanacearum* GM1000 in Arabidopsis wild-type (Col-0 WT) or *pub4* mutant plants. Surprisingly, *pub4* mutant plants showed weaker wilting symptoms than WT plants (Figures 4A, S3A, and S3B). To address the role of PUB4 in disease resistance in a natural host of *R. solanacearum*, we performed soil-drenching inoculation in tomato plants with roots either expressing a *SlPUB4* RNAi construct (*SlPUB4*-RNAi) or overexpressing *SlPUB4* (OE:*SlPUB4*). *SlPUB4*-RNAi plants showed a slight but reproducible reduction in wilting symptoms compared to control plants carrying an empty vector (EV-RNAi), demonstrating the same tendency as *pub4* mutants in Arabidopsis (Figures 4B and S3C-E). Accordingly, OE:*SlPUB4* plants showed earlier wilting symptoms than control EV plants, suggesting that *SlPUB4* overexpression promotes *R. solanacearum* infection (Figures 4C and S3F-H). Together, these results suggest that PUB4 acts as a positive regulator of *R. solanacearum* infection, and thus support the hypothesis that *PUB4* is a susceptibility gene for *R. solanacearum* that is targeted by RipAC.

**Figure 4.**
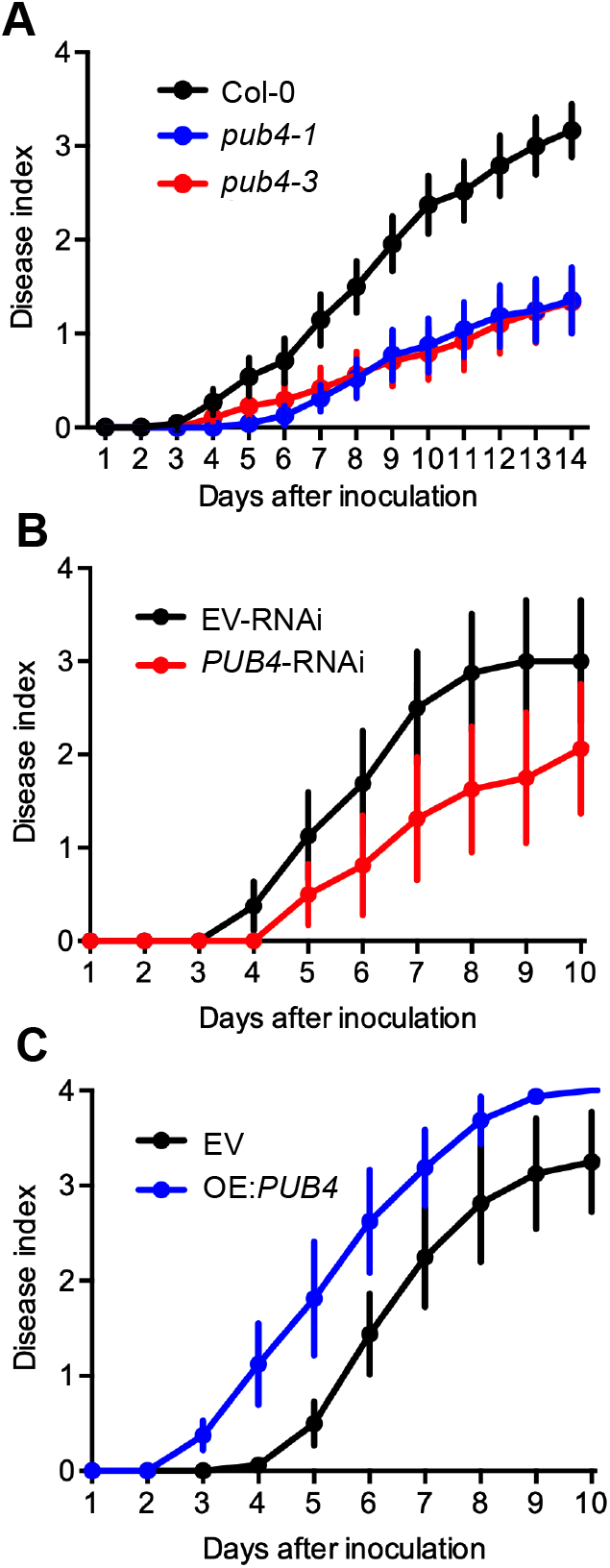
PUB4 promotes *R. solanacearum* infection in Arabidopsis and tomato. (A) Soil-drenching inoculation assays in Arabidopsis Col-0 and *pub4* mutant lines. Plants were inoculated with *R. solanacearum* GMI1000. The results are represented as disease progression, showing the average wilting symptoms in a scale from 0 to 4 (mean ± SE, n=24). (B-C) Soil-drenching inoculation assays in tomato plants with roots expressing a *PUB4* RNAi construct (*PUB4*-RNAi) (B) or overexpressing *PUB4* (OE:*PUB4*) (C). Plants were inoculated with *R. solanacearum* GMI1000. The results are represented as disease progression, showing the average wilting symptoms in a scale from 0 to 4 (mean ± SE, n=12). Experiments were performed at least 3 times with similar results. Panels show representative results; composite data from different experiments and survival analyses are shown in Figure S3.

### PUB4 associates with PRR complexes

To determine the molecular mechanism of PTI regulation by PUB4, we immuno-purified PUB4 and its interacting proteins from Arabidopsis transgenic plants expressing *PUB4-FLAG* either mock- or elf18-treated, and analysed the resulting immunoprecipitates by liquid chromatography followed by tandem mass spectrometry (LC-MS/MS). PUB4 was found to associate with the EFR-BAK1 PRR complex especially after elf18 treatment (Figure 5A). The extra-large G protein XLG2, known to associate with the FLS2 complex (Liang et al., 2016), and its close homologue XLG1, constitutively associated with PUB4 (Figure 5A). Surprisingly, we did not detect peptides of BIK1, which is known to associate with PRR complexes (Lu et al., 2010; Zhang et al., 2010). However, we have previously observed that the low protein accumulation of BIK1 can make it difficult to be detected in immunoprecipitates by LC-MS/MS. Thus, to determine whether PUB4 could also associate with BIK1, we co-expressed *BIK1-HA* with *PUB4-GFP* or *GFP* alone in *N. benthamiana*, and performed a co-immunoprecipitation (CoIP) before or after flg22 treatment. BIK1 was found to associate constitutively with PUB4 (Figure 5B). Moreover, a CoIP assay using *PUB4-FLAG* Arabidopsis plants further indicated that PUB4 associates with FLS2 and BAK1 specifically after flg22 treatment, and confirmed the constitutive association with BIK1 (Figure 5C).

**Figure 5.**
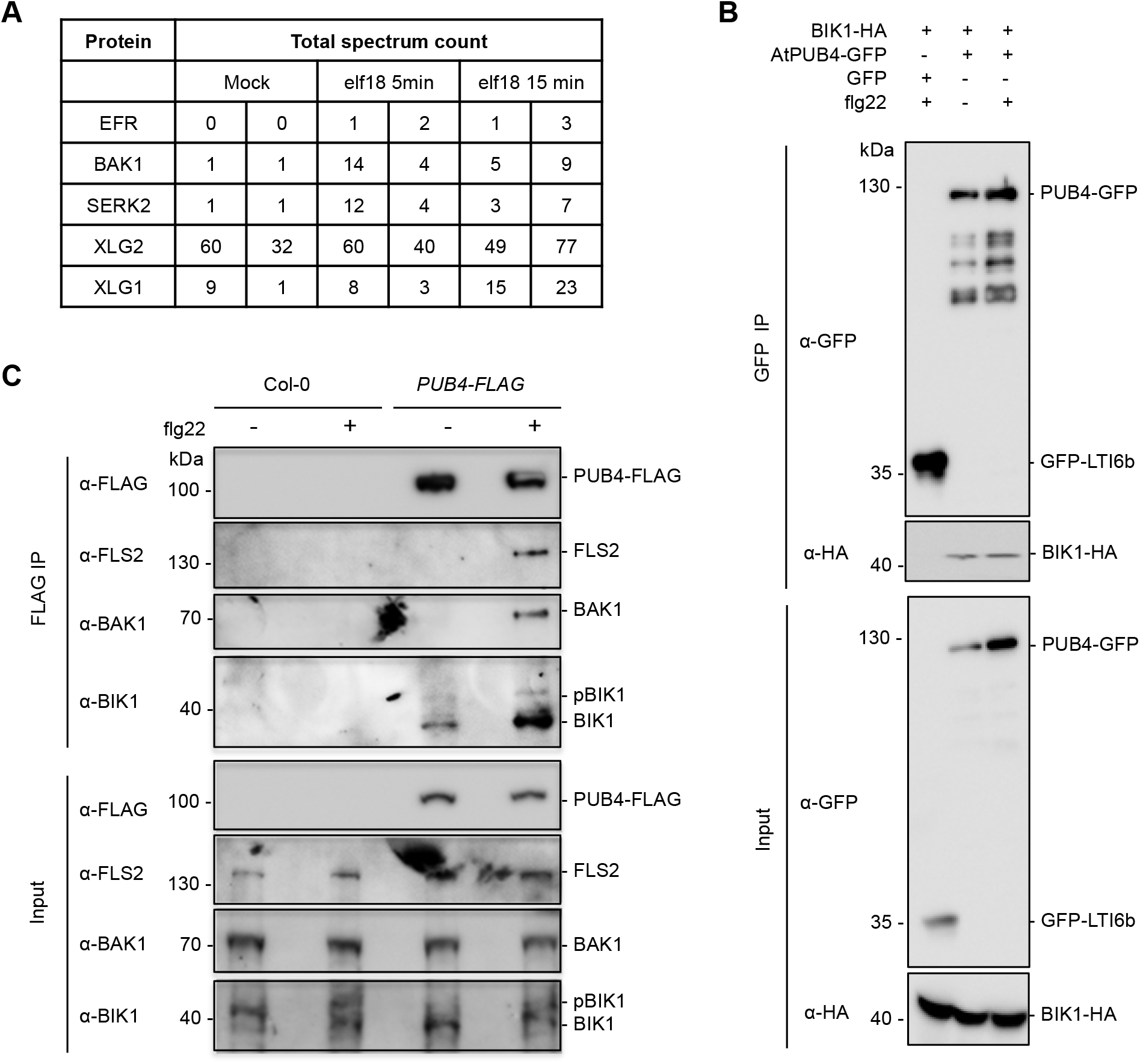
PUB4 associates with PRR complexes. (A) PUB4 associates in *Arabidopsis* with EFR, BAK1, and SERK2 after 1 mM elf18^Ecol^ treatment, and constitutively with XLG2 and XLG1. PRR complex members were identified after immunoprecipitation of PUB4-FLAG from *35S:PUB4-FLAG* line, tryptic digestion and sample analyses by LC-MS/MS. Untransformed Col-0 seedlings were used as a negative control. Total spectrum count for each protein is shown. (B) Co-immunoprecipitation of PUB4 and BIK1 in *N. benthamiana.* PUB4-GFP or GFP-LTI6b were transiently co-expressed with BIK1-HA in the leaves of *N. benthamiana.* After treatment with mock or 1 mM flg22^Paer^ for 10 min, total proteins (input) were extracted and subjected for immunoprecipitation with anti-GFP beads. Experiments were performed at least two times with similar results. (C) Co-immunoprecipitation of PUB4 with PRR complex members in *Arabidopsis*. PUB4 co-immunoprecipitated with FLS2 and BAK1 specifically after 10 min of 1 mM flg22^Paer^ treatment, and constantly with BIK1. Total protein extracts (input) from Col-0 and *PUB4-FLAG* plants were subjected to immunoprecipitation with anti-FLAG beads. Immunoblots were analyzed using the indicated antibodies. Molecular weight (kDa) marker bands are indicated for reference. Experiments were performed at least three times with similar results.

### RipAC does not affect PUB4 association with PRRs and BIK1

To understand the effect of RipAC on PUB4, we tested whether RipAC has an impact on PUB4 interaction with PRRs and BIK1. CoIP assays, using *PUB4-FLAG* or *PUB4-FLAG RipAC-GFP* plants, indicated that RipAC does not affect the constitutive association between PUB4 and BIK1, nor the flg22-dependent association between FLS2 and BAK1 (Figure 6). However, together with our previous results (Figure 5C), these assays revealed that PUB4 associates with modified BIK1, as we reproducibly observed a laddering on PUB4-associated BIK1 (Figure 6). Interestingly, this laddering was enhanced in the presence of RipAC (Figure 6). This observation led us to consider BIK1 as a relevant PUB4-associated protein and indirect RipAC target. Further support for this hypothesis comes from our genetic data demonstrating that both RipAC and PUB4 affect immune responses triggered by various PAMPs/DAMPs (Figures 1 and 3), and the known role of BIK1 as a convergent point downstream of multiple PRR complexes (Couto and Zipfel, 2016).

**Figure 6.**
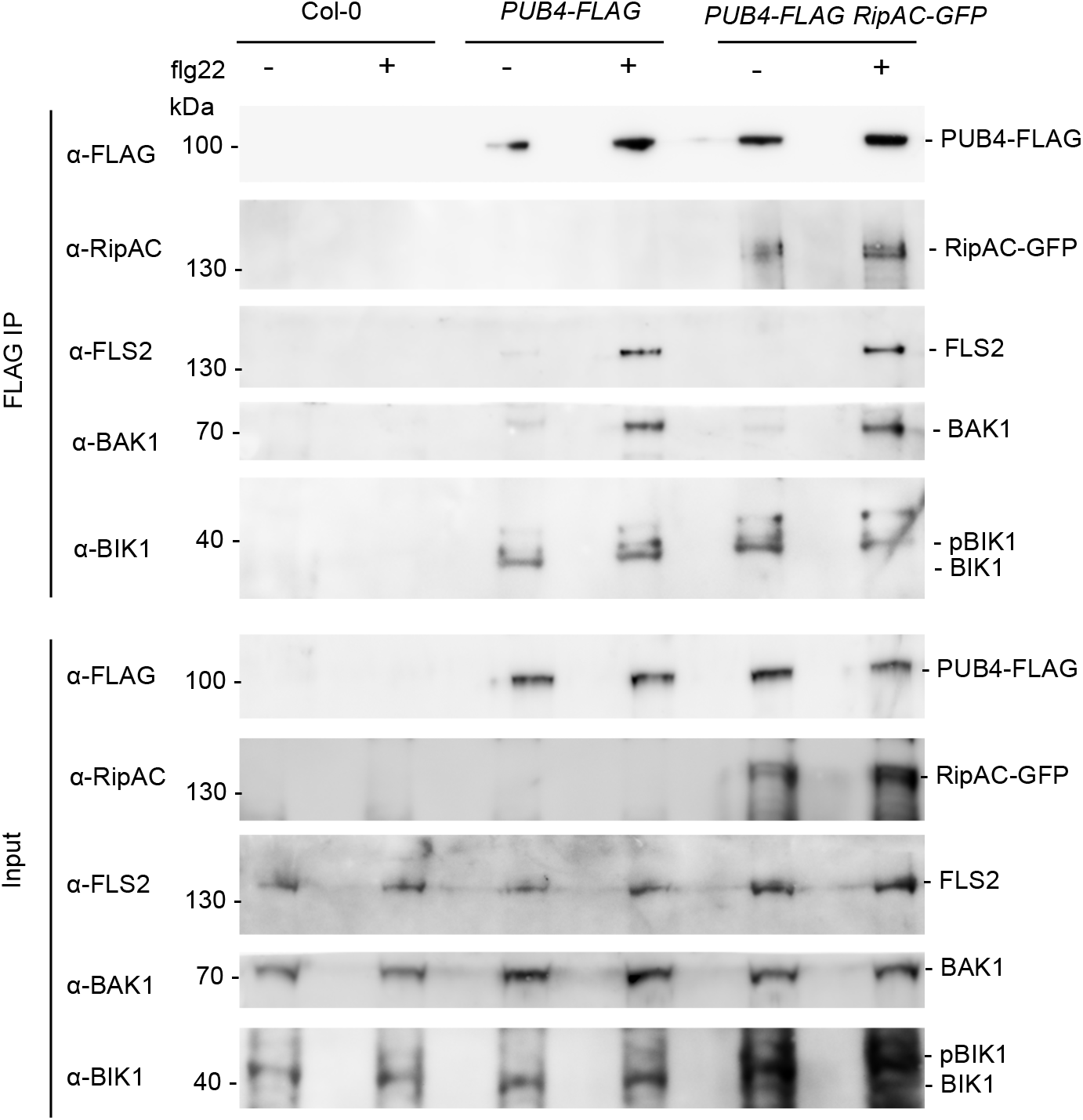
RipAC does not affect PUB4 accumulation or its association with PRRs and BIK1. RipAC does not affect PUB4 association with PRR complex members. *PUB4-FLAG, PUB4-FLAG RipAC-GFP* and Col-0 plants were treated for 10 min with water (as mock) or 1 mM flg22^Paer^ and elf18^Ecol^ treatment. Total protein extracts (input) were subjected for immunoprecipitation with anti-FLAG beads. Immunoblots were analyzed using the indicated antibodies. Molecular weight (kDa) marker bands are indicated for reference. The experiment was performed three times with similar results.

### PUB4 plays a dual role in the regulation of BIK1 protein homeostasis

To test whether PUB4 affects BIK1 protein levels, we used an anti-BIK1 antibody to analyse BIK1 accumulation in Arabidopsis WT and *pub4-1* mutant plants. We found that BIK1 accumulation was reduced in *pub4-1* plants after PAMP treatment (Figure 7A). To verify this result, we crossed the *BIK1-HA* line with the *pub4-1*^−^/^−^ mutant, but repeatedly failed to generate viable fertile *pub4* homozygous plants expressing *BIK1-HA*, suggestive of a genetic interaction between *PUB4* and *BIK1*. Thus, we analysed BIK1-HA accumulation in the *pub4-1*^+^/^−^ BIK1-HA^+^/^−^ F2 segregating population, using cycloheximide (CHX) treatment to inhibit *de novo* protein synthesis. The results show a reduced accumulation of BIK1 after flg22 treatment in *pub4-1*^−^/^−^ BIK1-HA^+^/^−^ plants compared to *BIK1-HA*^+^/^−^ plants (Figure 7B), suggesting that PUB4 function is required for accumulation of wild-type levels of activated BIK1. A lower mobility BIK1 band previously shown to correspond to phosphorylated BIK1 (Lu et al., 2010; Zhang et al., 2010) was detectable in *pub4-1*^−^/^−^ BIK1-HA^+^/^−^ plants, suggesting that the activation of BIK1 was not affected (Figure 7B). Inhibition of the proteasome activity using MG132 treatment abolished the reduction of BIK1 accumulation in the *pub4* mutant background (Figure 7B), suggesting that the reduced accumulation of activated BIK1 after treatment is due to its proteasomal degradation. The accumulation of non-activated BIK1 after CHX treatment was higher in *pub4-1*^−^/^−^ BIK1-HA^+^/^−^ plants compared to *BIK1-HA*^+^/^−^ (Figure 7B), and BIK1 phosphorylation in response to PAMP treatment was not affected in the *pub4-1* mutant background (Figure 7B), suggesting that a reduced amount of activated BIK1 in *pub4-1* is due to the inability of *pub4* mutant plants to preserve activated BIK1 from degradation. A requirement of PUB4 for the maintenance of signalling-competent BIK1 would explain the impaired immune responses in *pub4* mutant plants in response to various PAMPs/DAMPs.

**Figure 7.**
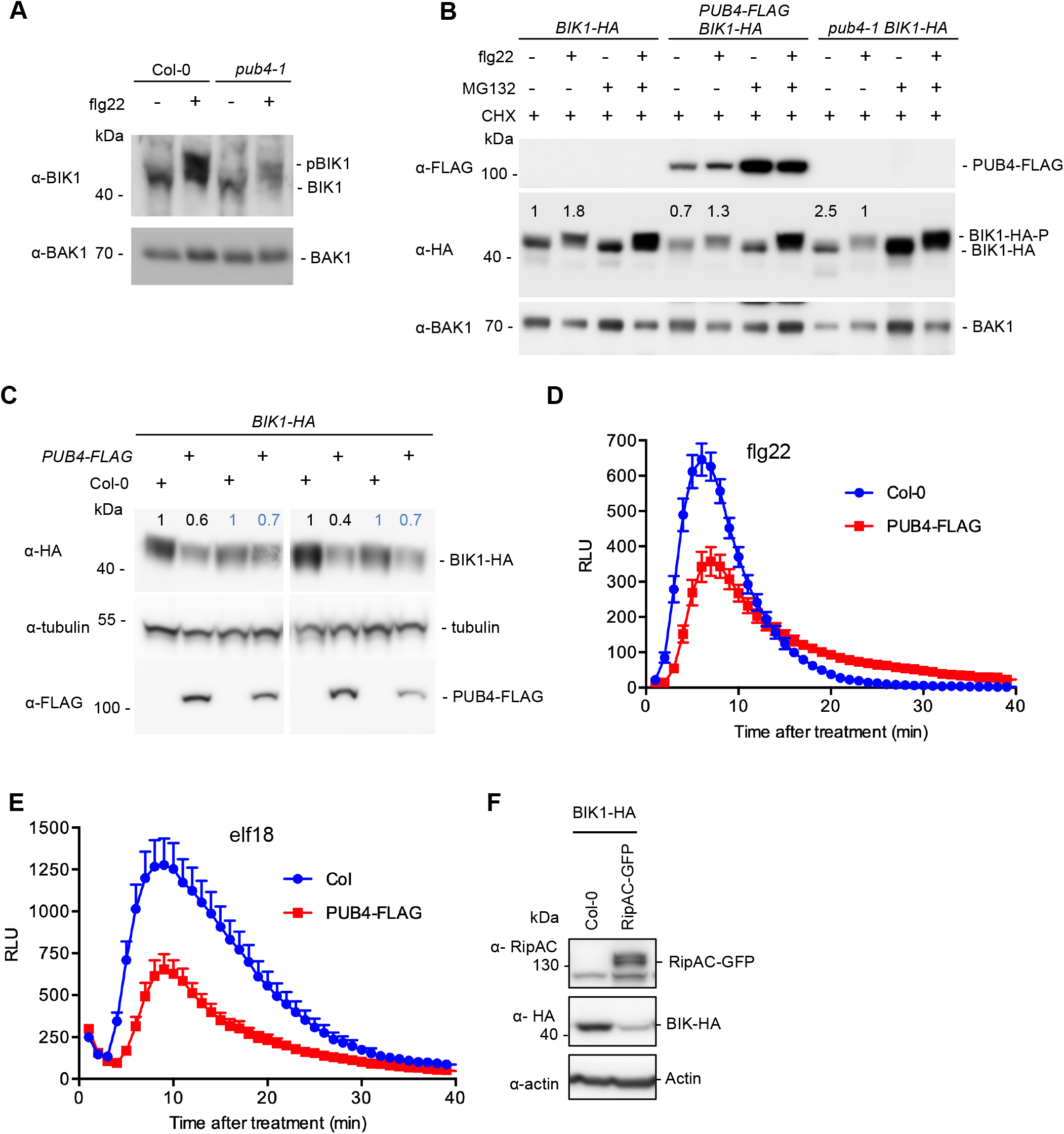
BIK1 accumulation is dually regulated by PUB4 and negatively affected by RipAC. (A) Analysis of BIK1 protein accumulation in Col-0 and *pub4* seedlings with or without treatment with 1 mM flg22^Paer^ for 10 min. BAK1 was used as a loading control. Protein extracts were enriched for membrane-associated proteins. Experiments were performed at least 3 times with similar results. (B) Analysis of BIK1-HA protein accumulation in *BIK1-HA*^+^/^−^, *pub4*^−^/^−^ *BIK1-HA*^+^/^−^, and *PUB4-FLAG*^+^/^−^ *BIK1-HA*^+^/^−^ in 4- to 5-week-old plants. Prior to protein extraction leave disks were treated for 6 h with 100 mM MG132, 50 mM CHX, and for 10 min with water (as mock) or 1 mM flg22^Paer^. Proteins were extracted in SDS buffer and analysed by western blot. BAK1 was used as a loading control. Experiments were performed at least 3 times with similar results. (C) BIK1-HA accumulation is reduced in *PUB4-FLAG*^+^/^−^ *BIK1-HA*^+^/^−^ line compared to *BIK1-HA*^+^/^−^ line in the absence of PAMP treatment. Total protein extracts were analysed by western blot. Tubulin was used as a loading control. (D-E) PUB4 overexpression leads to reduction in ROS burst in response to (D) 100 nM flg22^Paer^ and (E) 100 nM elf18^Ecol^. ROS was measured as relative luminescence units (RLU) over time. Values are means ± SE (n=24). Experiments were performed at least 3 times with similar results. (F) BIK1-HA accumulation is reduced in *RipAC-GFP*/*BIK1-HA* line compared to Col-0/*BIK1-HA* line. Total protein extracts were analysed by western blot. Actin was used as a loading control. In (B), (C), and (F), WB quantification was performed using ImageJ software. Immunoblots were analyzed using the indicated antibodies. Molecular weight (kDa) marker bands are indicated for reference. Experiments were performed at least three times with similar results.

Interestingly, we found lower accumulation of non-activated BIK1 in *PUB4-FLAG*^+^/^−^ *BIK1-HA*^+^/^−^ plants, which overexpress *PUB4*, compared to *BIK1-HA*^+^/^−^ plants (Figures 7B, 7C, and S4A). This is consistent with the higher accumulation of non-activated BIK1 observed in *pub4* plants (Figure 7B). BIK1 activation, detected by the presence of lower mobility (phosphorylated) BIK1, was not affected in *PUB4* overexpression line (Figure 7B). Several observations suggest that PUB4 promotes the degradation of non-activated BIK1: (*i*) PUB4 associates with BIK1 in basal conditions, and there is a laddering on PUB4-associated BIK1; (*ii*) *PUB4* overexpression leads to reduced accumulation of non-activated BIK1; and (*iii*) *pub4* mutation leads to enhanced accumulation of non-activated BIK1. In accordance with this hypothesis, and considering that BIK1 is a rate-limiting factor in the activation of early PTI responses, PAMP-triggered ROS production was impaired in the *PUB4-FLAG* overexpressing line (Figures 7D, E). Thus, PUB4 seems to play a dual role in BIK1 stability: promoting the degradation of non-activated BIK1, while preserving activated BIK1 from degradation.

### RipAC negatively impacts BIK1 accumulation

As PUB4 regulates BIK1 stability and is targeted by RipAC, we tested whether RipAC affects BIK1 protein accumulation. For this, we crossed *RipAC-GFP* plants with *BIK1-HA* plants, and analysed the resulting F1. Plants expressing *RipAC* showed lower BIK1 accumulation compared to control plants (Figures 7F and S4B), which could explain the impaired ROS burst in these plants in response to various PAMPs. Despite the impact of RipAC on BIK1 accumulation, RipAC did not associate with BIK1 in CoIP and Split-LUC assays *in planta* (Figure S5). This suggests that RipAC acts on BIK1 via PUB4, which is supported by the greater band shift of PUB4-associated BIK1 observed after flg22 treatment in the *PUB4-FLAG RipAC-GFP* line compared to *PUB4-FLAG* plants (Figure 6).

### RipAC causes a reduction in PUB4 accumulation after PAMP treatment and manipulates PUB4 phosphorylation

The fact that PUB4 dually regulates BIK1 suggests that there are two pools of PUB4: ‘pre-elicitation’ and ‘post-elicitation’, which affect BIK1 differently. PUB4 has been previously found among proteins rapidly phosphorylated in response to PAMP treatment (Benschop et al., 2007), supporting this idea. Thus, we hypothesized that RipAC might shift the balance between the two forms of PUB4 in favour of the ‘pre-elicitation’ state to promote BIK1 degradation, possibly through the control of PUB4 protein accumulation or posttranslational modifications. To test this hypothesis, we purified PUB4 from *PUB4-FLAG* or *PUB4-FLAG RipAC-GFP* plants, either mock- or PAMP-treated, and analysed the samples by LC-MS/MS and parallel reaction monitoring (PRM) LC-MS/MS. Notably, PUB4 accumulation increased after flg22 and efl18 treatment, and this effect was abolished by RipAC (Figure 8A). This suggests that RipAC diminishes the positive role of PUB4 in the stabilization of activated BIK1 and therefore promotion of PTI.

**Figure 8.**
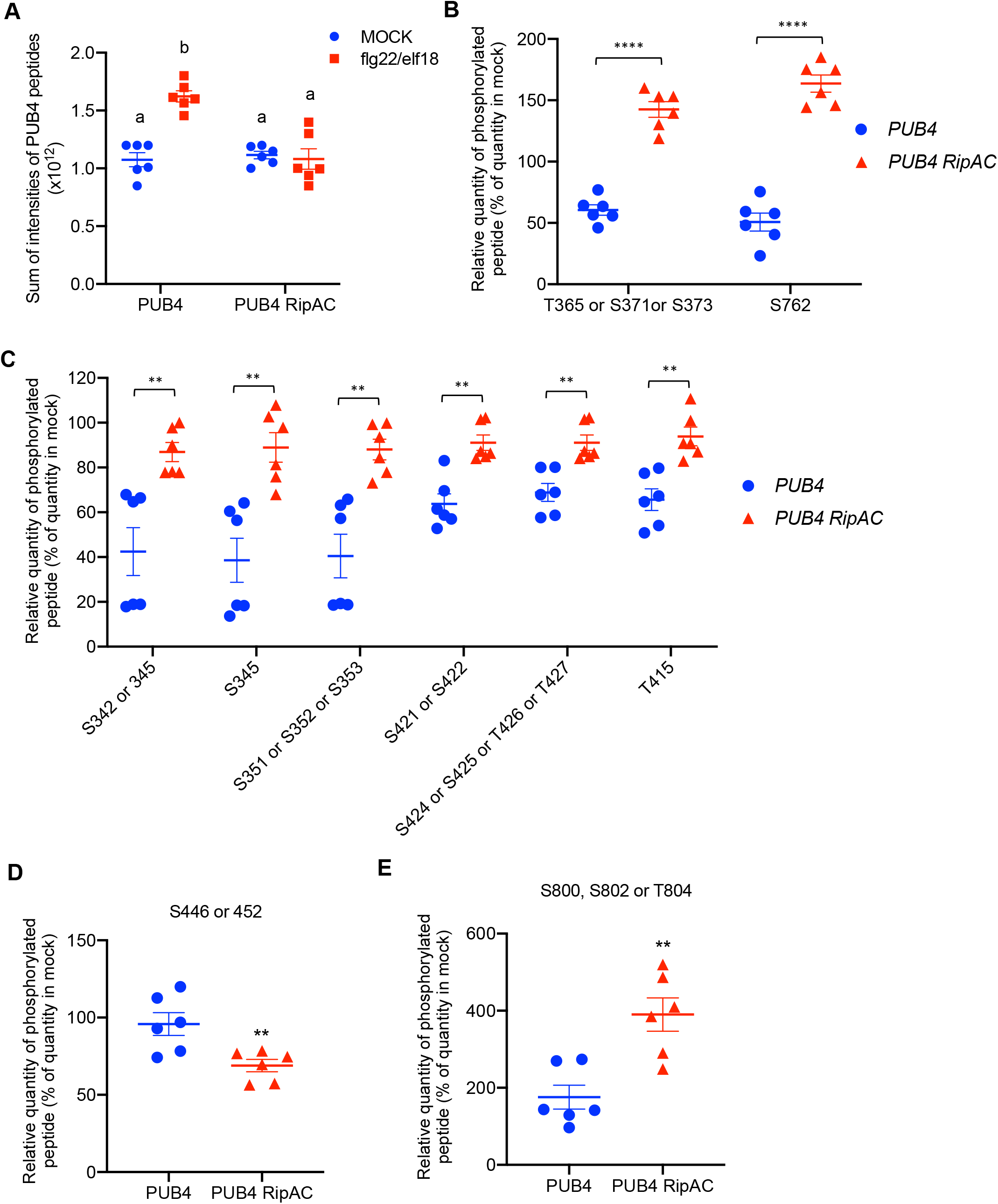
RipAC causes a reduction in PUB4 accumulation after PAMP treatment and manipulates PUB4 phosphorylation. (A) RipAC prevents an increase in PUB4 accumulation after PAMP treatment. PUB4 was immunoprecipitated from *PUB4-FLAG* or *PUB4-FLAG RipAC-GFP* plants after treatment with PAMPs (1 mM flg22^Paer^, 1 mM elf18^Ecol^) or water (as mock treatment). The samples were digested with trypsin and analysed by parallel reaction monitoring (PRM) LC-MS/MS. Values represent the sum of intensities of PUB4 corresponding peptides. Data are means ± SE of two biological replicates, each of which contains three technical replicates (two-way ANOVA, Tukey’s multiple comparison test). Different letters indicate significantly different values at p<0.0001). (B-E) RipAC affects PUB4 phosphorylation after PAMP treatment. The samples were prepared and processed as in (A). The abundance of phosphorylated peptide was calculated as the ratio of intensities of phosphorylated vs PUB4 control peptide sum. Values represent the percentage of PUB4 phosphorylated peptide in PAMP-treated vs mock-treated seedlings, with 100 % indicating the same level of PUB4 phosphorylation in PAMP- and mock-treated seedlings. Values are means ± SE of two biological replicates, each of which contains three technical replicates (t test, asterisks indicate significant differences compared to *PUB4-FLAG*, ****p<0.0001, **p<0.01).

Our preliminary data showed that PUB4 is a highly phosphorylated protein both before and after PAMP treatment. We thus quantified PUB4 phosphorylation by PRM LC-MS/MS in the absence and presence of RipAC, with either mock or PAMP treatment. As effectors are typically delivered once PTI has already been activated, we focused on analyses of PUB4 phosphosites affected by RipAC only after PAMP but not mock treatment. Thus, we identified PUB4 phosphosites manipulated by RipAC and showed percent of their relative phosphorylation after PAMP treatment compared to mock treatment (Figures 8B-E). Most PUB4 phosphosites affected by RipAC show the same tendency: PUB4 phosphosites that were significantly downregulated upon PAMP treatment are attenuated or upregulated in the presence of RipAC (Figure 8B and 8C), with two PUB4 regions demonstrating different tendencies (Figure 8D-E). Most of the affected phosphosites lie in the hinge region of PUB4, between the U-box and ARM repeats (Figure S1A), while S762 is in the ARM domain responsible for protein-protein interaction. Hence, S762 phosphorylation status may affect PUB4 interaction with other proteins. These data suggest that RipAC impairs the decreased phosphorylation on these sites after PAMP treatment, thus promoting the ‘pre-elicitation’ state of PUB4, in which PUB4 promotes BIK1 degradation.

## Discussion

Immune signalling is tightly regulated to prevent autoimmunity while ensuring optimal strength and duration of immune responses upon detection of potential pathogens. This is achieved through negative regulators acting at the level of PRR complex formation and activation, cytoplasmic signal transduction, and activation of defence-related genes (Couto and Zipfel, 2016). Although this provides a robust mechanism for the regulation of immune activation, it also constitutes a potential target for manipulation by invading pathogens.

In this study, we identified the U-box E3 ubiquitin ligase PUB4 as a target of the *R. solanacearum* effector RipAC. Detailed genetic analysis indicated that PUB4 is a common regulator of immune signalling triggered by various PAMPs/DAMPs. We show that PUB4 is recruited to FLS2/EFR-BAK1 complexes after PAMP treatment, and constitutively associates with BIK1 and XLG1/XLG2. Our biochemical data is consistent with previous reports showing that PUB4 associates with XLG1/2/3 *in planta* (Wang et al., 2017), and that XLG2 associates with PRR complexes in basal conditions (Liang et al., 2016). Interestingly, we did not detect association of PUB4 with FLS2 before PAMP treatment, and other heteromeric G protein complex subunits were not detected among PUB4-associated proteins, suggesting that PUB4 associates with activated (GTP-bound) XLG2 and BIK1 pools not associated with FLS2. A recent report showed that PUB4 associates with CERK1, and is a positive regulator of chitin-triggered signalling (Desaki et al., 2019). Our data showing that PUB4 not only positively regulates chitin responses, but also flg22, elf18 and AtPep1 responses, together with the known role of BIK1 downstream of their respective PRRs, suggest that the association of PUB4 and BIK1 actually explains the genetic contribution of PUB4 to responses triggered by diverse elicitors.

Our results also provide a mechanism explaining the effect of PUB4 on immune responses triggered by various PAMPs/DAMPs, since we show that PUB4 regulates BIK1 protein homeostasis. Being a hub of immune signalling and a rate-limiting factor in PTI responses, BIK1 accumulation and activation are tightly regulated (Couto et al., 2016; Kadota et al., 2014; Liang et al., 2016; Monaghan et al., 2014; Wang et al., 2018; Zhang et al., 2010; Zhang et al., 2018). The current model of BIK1 regulation postulates the existence of two BIK1 pools: ‘non-activated’ (before PAMP treatment) and ‘activated’ (after PAMP treatment). Furthermore, non-activated BIK1 can be associated or not to PRR complexes. CPK28 does not associate with FLS2, but constantly associates with BIK1 (Monaghan et al., 2014). In this scenario, PRR-associated BIK1 is protected from degradation by PRR-associated G protein complexes, while free BIK1 is preferentially phosphorylated by CPK28 and subsequently targeted for degradation by the E3 ubiquitin ligases PUB25/26 (Wang et al., 2018). Notably, BIK1 accumulation is higher in *cpk28* mutant plants compared to *pub25 pub26* mutant plants, suggesting that other E3 ubiquitin ligase(s) are involved in BIK1 degradation (Wang et al., 2018). In this study, we found that PUB4 promotes degradation of non-activated BIK1, which correlates with the fact that PUB4 associates with BIK1, but not with FLS2, in basal conditions. It is noteworthy that PUB25/26 are negative regulators of immune activation, promoting the degradation of non-activated BIK1, while PUB4 has a similar effect before PAMP treatment, but promotes the accumulation of active BIK1 after PAMP treatment. Plants that over-express PUB4 would accumulate less BIK1 before PAMP treatment; therefore, rapid immune responses such as the burst of ROS would be affected, reflecting the effect of PUB4 as a negative regulator before PAMP treatment. However, *pub4* mutants show a lack of activated BIK1, which explains why immune responses are down-regulated, demonstrating that, after PAMP treatment, PUB4 indeed acts as a positive regulator of immunity. This represents two distinct modes of BIK1 regulation executed by PUB4 and PUB25/26.

We found that PUB4 is required for the accumulation of wild-type levels of activated BIK1, which explains the reduction of PTI responses observed in *pub4* mutant plants. Notably, BIK1 phosphorylation in response to PAMP treatment is not affected in *pub4* plants, suggesting that PUB4 prevents the degradation of activated BIK1. The dual role of PUB4 in the regulation of BIK1 could be explained by the existence of two pools of PUB4: ‘pre-elicitation’ and ‘post-elicitation’. In accordance with the fact that PUB4 was previously found to be rapidly phosphorylated after PAMP treatment (Benschop et al., 2007), we demonstrated that PUB4 is a highly phosphorylated protein, and that its phosphorylation status correlates with its function in immunity. Moreover, PAMP treatment leads to an increase in PUB4 accumulation, which promotes its role in the stabilization of activated BIK1. While we reveal here that PUB4 positively regulates activated BIK1 accumulation, the E3 ubiquitin ligases RING-H2 FINGER A3A (RHA3A) and RHA3B were recently shown to promote BIK1 activation (Ma et al., 2020); thus illustrating that distinct BIK1 ubiquitination events positively regulate both its accumulation and activation.

Supporting its important role in the regulation of plant immunity, we found that PUB4 is targeted by the *R. solanacearum* T3E RipAC. RipAC does not disrupt PUB4 association with FLS2-BAK1-BIK1 complex, but causes a reduction in PUB4 accumulation after PAMP treatment, thus diminishing PUB4 ‘post-elicitation’ pool that positively regulates BIK1 and PTI. Moreover, RipAC manipulates PUB4 phosphorylation state with the main tendency to promote phosphorylation of ‘pre-elicitation’ PUB4. We found that RipAC dramatically reduces BIK1 accumulation without interacting with BIK1. Considering the effect of PUB4 on the accumulation of non-activated BIK1, our results suggest that RipAC exploits PUB4 and promotes accumulation of the ‘pre-elicitation’ form of PUB4 to reduce BIK1 accumulation. Interestingly, although *pub4* mutants were more susceptible to non-pathogenic *P. syringae*, PUB4 was found to promote disease symptoms caused by *R. solanacearum* infection. The opposite results observed in the genetic analysis of the role of PUB4 in resistance against *Pto* and *R. solanacearum* could be due to specific requirements for PUB4 in various plant tissues. This seems however unlikely considering that *pub4* mutants also show reduced flg22-triggered ROS in root tissues (Figure S6), suggesting a positive role of PUB4 in the regulation of early PTI responses also in roots. However, lack of PUB4 in leaves leads to inability of plants to close stomata in response to pathogen attack, which is an important factor contributing to virulence of *Pto* DC3000 COR^*−*^ and *Pto* DC3000 *hrcC*. *pub4* plants have also recently been shown to exhibit elevated levels of the defence-related hormone salicylic acid (SA), and enhanced expression of defence genes (Desaki et al., 2019). Although one would expect such SA disbalance to affect similarly both *Pto* and *R. solanacearum*, which are both hemi-biotrophic pathogens, we cannot fully exclude the possibility that *R. solanacearum* and *Pto* are affected differently by increased SA levels, especially considering the different tissues infected by these bacteria. Our data show that RipAC dually affects PUB4: by attenuating its positive role in immunity and by promoting its negative role in accumulation of non-activated BIK1. The latter makes PUB4 an important susceptibility factor required for optimal *R. solanacearum* infection. This could explain why PUB4 promotes *R. solanacearum* infection, despite playing a positive role in PTI and resistance against *Pto*.

Future work should determine the molecular mechanisms by which PUB4 phosphorylation affects PUB4 interaction with other proteins and its E3 ubiquitin ligase activity during immunity.

## Acknowledgments

This project has received funding from the Strategic Priority Research Program of the Chinese Academy of Sciences (grant XDB27040204 to A.P.M.), the National Natural Science Foundation of China (grant 31571973 to A.P.M.), the Chinese 1000 Talents Program (to A.P.M.), the Shanghai Center for Plant Stress Biology (to A.P.M.), the China Postdoctoral Science Foundation (fellowship 2016M600339 to G.Y.), the President’s International Fellowship Initiative (PIFI) (fellowships 2018PB0057 and 2020PB0088 to J.S.R.), the European Union’s Horizon 2020 research and innovation programme under the Marie Skłodowska-Curie grant agreement No 753641 (to M.D.), the Gatsby Charitable Foundation (to C.Z.), the European Research Council under the Grant Agreement No 309858 (grant “PHOSPHinnATE” to C.Z.), the University of Zürich (to C.Z.), and the Swiss National Science Foundation (grant 31003A_182625) (to C.Z.). S.J. was supported by a post-doctoral fellowship from the European Molecular Biology Organization (EMBO-LTF #225-2015). We thank Prof. Nick Talbot and his group for hosting M.D. and fruitful discussions. We thank Prof. Yiji Xia and Prof. Jian-Min Zhou for sharing biological materials. We thank Matthew Smoker, Jodie Taylor and Juan Lopez from the TSL Plant Transformation support group for plant transformation, the John Innes Centre Horticultural Services for plant care, the PSC Cell Biology core facility for assistance with confocal microscopy, Xinyu Jian for technical and administrative assistance, and all past and current members of the Zipfel and Macho groups for technical help and fruitful discussions.

## Author Contributions

A.P.M. and C.Z. supervised the project and obtained the funding. M.D., G.Y and J.S.R. conceived, designed, and performed the majority of plant and biochemical experiments and obtained the funding. S.J. provided initial data for the project, performed seedling growth inhibition assay, and a CoIP in *N. benthamiana*. P.D. and F.M. performed mass-spectrometry analyses. R.J.L.M. performed root transformation and subsequent bacterial inoculation assays. L.S. assisted with molecular cloning. Y.W. performed ROS assays in roots. M.D, C.Z., and A.P.M. wrote the manuscript. All authors commented and agreed on the manuscript before submission.

## Materials and Methods

### Resources

All the biological and chemical materials used in this study are summarized in Table S1.

#### Arabidopsis thaliana

*Arabidopsis thaliana* materials in this study are derived from ecotype Columbia (Col-0). Previously published lines include: *NP::BIK1-HA* (Zhang et al., 2010), *35S::PUB4-FLAG* (Wang et al., 2013), RipAC transgenic lines (RipAC #3 and RipAC #31) (Yu et al., 2020). The T-DNA insertion lines *pub4-1* (SALK_108269) and *pub4-3* (SAIL_859_H05) (Wang et al., 2013) were obtained from the Nottingham Arabidopsis Stock Centre (NASC), and homozygous lines were selected by genotyping using allele-specific primers.

In the experiments with Arabidopsis seedlings in 1/2 MS media, the seedlings were kept on 1/2 MS plates in a growth chamber (22 °C, 16 h light/8 h dark, 100-150 mE m^−2^ s^−1^) for germination and growth for 5 days, then transferred to 1/2 MS liquid culture for additional 7-9 days. For PAMP-triggered ROS burst assays, *Pseudomonas syringae* and *Ralstonia solanacearum* infection assays, Arabidopsis plants were grown in either soil or jiffy pots (Jiffy International, Kristiansand, Norway) in a short day chamber (22 °C, 10 h light/14 h dark photoperiod, 100-150 mE m^−2^ s^−1^, 65 % humidity) for 4-5 weeks. After soil drenching inoculation, the plants were transferred to a growth chamber controlled with the following conditions: 75 % humidity, 12 h light, 130 mE m^−2^ s^−1^, 27 °C, and 12 h darkness, 26 °C for disease symptom scoring.

#### Nicotiana benthamiana

*Nicotiana benthamiana* plants were cultivated at 22 °C in a walk-in chamber under 16 h light/8 h dark cycle and a light intensity of 100-150 mE m^−2^ s^−1^.

#### Solanum lycopersicum

Tomato plants (*Solanum lycopersicum* cv. Moneymaker) were cultivated in jiffy pots (Jiffy International, Kristiansand, Norway) in controlled growth chambers (25 °C, 16 h light/8 h dark photoperiod, 130 mE m^−2^ s^−1^, 65% humidity) for 4 weeks. After soil drenching inoculation, the plants were kept in a growth chamber under the following conditions: 75% humidity, 12 h light, 130 mE m^−2^ s^−1^, 27 °C, and 12 h darkness, 26 °C for disease symptom scoring. To grow tomato seedlings *in vitro*, tomato seeds were surface-sterilized by soaking in 5% (v/v) sodium hypochlorite for 5 minutes, washed 4-5 times with distilled sterile water and shaken slowly in sterile water overnight to facilitate germination. Then, seeds were germinated on half-strength of Murashige and Skoog medium without sucrose (2.21 g/L MS, 0.8% w/v agar) for 3-4 days at 25 °C, in darkness.

### Bacterial strains

*Pseudomonas syringae* pv. *tomato* (*Pto*) DC3000 strains, including *Pto* containing an empty vector (EV), or the type-three secretion system (T3SS)-defective mutant *ΔhrcC*, or coronatine-defective mutant *COR*^−^ were cultured overnight at 28 °C in LB medium containing 25 ◻g mL^−1^ rifampicin, and 25 ◻g mL^−1^ kanamycin.

*Ralstonia solanacearum* GMI1000 wild-type strain was grown overnight at 28 °C in complete BG liquid medium (Plener et al., 2012).

*Agrobacterium tumefaciens* GV3101 and *Agrobacterium rhizogenes* MSU440 with different constructs were cultured grown at 28 °C on LB agar media with appropriate antibiotics. The concentration for each antibiotic is: 25 ◻g mL^−1^ rifampicin, 50 ◻g mL^−1^ gentamicin, 50 ◻g mL^−1^ kanamycin, 50 ◻g mL^−1^ spectinomycin.

### Constructs and transgenic plants

The primers used to generate constructs in this work are listed in Table S2.

To generate constructs for co-immunoprecipitation and split-luciferase complementation (Split-LUC) assays, coding sequences were amplified by PCR using cDNA as template and inserted into corresponding destination vectors by either gateway cloning, golden-gate cloning system or In-fusion cloning. The recombinant construct was transformed into *A. tumefaciens* GV3101.

Arabidopsis lines generated in this study include: *35S::RipAC-GFP*/*NP::BIK1-HA*, *35S::RipAC-GFP*/*35S::PUB4-FLAG*, *35S::PUB4-FLAG/NP::BIK1-HA*, *pub4-1/NP::BIK1-HA.* These lines were generated by crossing, and confirmed by allele-specific primers, antibiotics screening and/or western blotting.

#### *Pseudomonas syringae* infection assays

For *Pto* inoculation, different *Pto* strains were resuspended in water at OD_600_=0.1 (5×10^7^ CFU mL^−1^). Before spraying, final concentration of 0.02 % Silwett L-77 was added to the inoculum. The bacterial suspensions were sprayed on 3- to 4-week-old Arabidopsis leaves. Bacterial numbers were determined 3 days post-inoculation (dpi). The whole plants were harvested in Eppendorf tubes and weighed.

#### *Ralstonia solanacearum* infection assays

For *R. solanacearum* soil drenching inoculation, 15 four-to-five-week-old Arabidopsis plants per genotype or 12 tomato plants (grown in Jiffy pots) were inoculated by soil drenching with a bacterial suspension containing 10^8^ colony-forming units per mL (CFU mL^−1^) as previously described (Yu et al., 2020). Briefly, 300 mL of inoculum of GMI1000 strain was poured to soak each treatment. Plants were transferred from the bacterial solution to a bed of potting mixture soil in a new tray (Vailleau et al., 2007) after 20-min incubation with the bacterial inoculum. To score the visual disease symptoms, a scale ranging from ‘0’ (no symptoms) to ‘4’ (complete wilting) was performed as previously described (Vailleau et al., 2007).

### ROS production assays

ROS measurements were performed in Arabidopsis plants as described previously (Sang and Macho, 2017). Plant leaf discs were collected and floated on sterile water overnight in 96-well plate. The water was then removed and replaced with 100 μL of eliciting solution containing 17 mg/mL luminol (Sigma Aldrich), 200 μg/mL horseradish peroxidase (Sigma Aldrich), and an appropriate concentration of the desired PAMP: 50 nM flg22^Pto^, 100-200 μg mL^−1^ chitin, 100 nM elf18^Pto^, 100 nM elf18^Rsol^, 100 nM flg22^Paer^ or 100 nM elf18^Ecol^. ROS assays in roots were performed as previously described (Wei et al., 2018). Briefly, Arabidopsis seeds were first germinated on 1/2 MS solid medium for 7 days and then transferred to 1/2 MS liquid culture for 5 days before ROS measurement. Roots of seedlings were cut into one-centimetre-long sections and allowed to recover for 5 h in 96-well plates with 100 μL H_2_O in each well. Sixteen root sections were analysed for each sample, using 100 nM of flg22^Pto^. For root assays, luminol was replaced by the more sensitive derivative L-012 (Wako Chemical, Japan). The luminescence was measured over 60 min using a Microplate luminescence reader (Varioskan flash, Thermo Scientific, USA) or a charge-coupled device camera (Photek Ltd., East Sussex UK).

### Seedling growth inhibition

Sterile Arabidopsis seeds were sown on MS 1% sucrose agar plates. The seeds were stratified in the dark at 4°C for 3-4 days and then transferred to light (LD). Four days later, one seedling per well was transferred to a 48-well plate containing 500 μl of sterile liquid MS 1 % sucrose supplemented either with water (as mock treatment) or PAMP (100 nM flg22 or 100 nM elf18). 12 seedlings per condition were used. Seedlings were transferred back to light for 10 days. Fresh weight of each seedling after blotting dry was recorded.

### Stomatal closure assays

Leaf discs (two leaf discs per plant, four plants per line) were taken from 4-to 5-week-old plants grown on soil and incubated in stomatal opening buffer (10 mM MES-KOH, pH 6.1; 50 mM KCl; 10 μM CaCl2; 0.01 % Tween-20) for 2 h in a plant growth cabinet in the light. Subsequently, 10 μM flg22 or mock; 1 mg/mL chitin or mock; 10 μM ABA or mock were added, and samples were incubated under the same conditions for another 2-3 h. Photographs of the abaxial leaf surface were taken using a Leica DM5500 microscope equipped with a Leica DFC450 camera. Width and length of the stomatal openings were determined using Image J software and stomatal aperture is shown as ratio of width divided by length. Values are means ± SE (n>120). Samples were analysed by one-way ANOVA, Kruskal-Wallis test and Dunn’s multiple comparisons test.

### Transient expression in *N. benthamiana*

For Split-LUC, FRET-FLIM, and co-immunoprecipitation assays, *A. tumefaciens* GV3101 or AGL1 (for epiGreen-*35S::PUB4-GFP*) carrying desired constructs were infiltrated into leaves of 5-week-old *N. benthamiana.* The OD600 used was 0.5 for each strain in all the assays, except for Split-LUC assays, for which we used OD600=0.2. *A. tumefaciens* was incubated in the infiltration buffer (10 mM MgCl_2_, 10 mM MES pH 5.6, and 150 μM acetosyringone) at room temperature for 2 h. Samples were collected 2 or 3 d after infiltration.

### Protein extraction and western blot assays

For protein extraction in plant tissues, 12 fourteen-day-old Arabidopsis seedlings or leaf discs (diameter=18mm) from *N. benthamiana* were frozen in liquid nitrogen and ground with a Tissue Lyser (QIAGEN, Hilden, Nordrhein-Westfalen, Germany). Protein samples were extracted with buffer containing 100 mM Tris (pH 8), 150 mM NaCl, 10% Glycerol, 1% IGEPAL, (5 mM EDTA, optional), 5 mM DTT, 1% Protease inhibitor cocktail, 2 mM PMSF, 10 mM sodium molybdate, 10 mM sodium fluoride, 2 mM sodium orthovanadate. The resulting protein samples were boiled at 70 °C for 10 min in Laemmli buffer and loaded in SDS-PAGE acrylamide gels for western blot. Alternatively, proteins from leave disks of 4-to 5-week-old Arabidopsis plants were extracted by adding 2x SDS buffer and heating at 70 °C for 10 min. Western blot detection was done using either PierceTM ECL western blotting substrate or SuperSignal™ West Femto Maximum Sensitivity Substrate (ThermoFisher). All the immunoblots were probed using appropriate antibodies as indicated in the figures. Molecular weight (kDa) marker bands are indicated for reference. Western blot quantification was performed using ImageJ software (http://imagej.net/). For enrichment of membrane associated proteins MinuteTM Plasma Membrane Protein Isolation Kit for Plants (Invent Biotechnologies) was used.

### MAPK activation assays

PAMP-triggered MAPK activation was evaluated as previously described (Macho et al., 2012). Briefly, 2 twelve-day-old Arabidopsis seedlings were treated with 100 nM flg22 and samples were collected at different time points. After protein extraction, the protein samples were loaded in 10% SDS-PAGE gels and the western blots were analyzed with anti-pMAPK antibodies. Blots were stained with Coomassie Brilliant Blue (CBB) to verify equal loading.

### Co-immunoprecipitation

Co-immunoprecipitation assays were performed as previously described (Kadota et al., 2016; Sang et al., 2018), with some changes. One gram of *N. benthamiana* leaf tissues was collected at 2 days after agro-infiltration and frozen in liquid nitrogen. Sterilized Arabidopsis seeds were grown on MS 1% sucrose agar plates for one week. Then, the seedlings were transferred into 6-well plates containing liquid MS; 5 seedlings per well. Two-week-old seedlings from two 6-well plates were treated by elicitor (1 μM elf18 or/and flg22) or MS medium (as mock treatment) for 10 min, including 2 min vacuum infiltration. Seedlings were frozen and then ground in liquid nitrogen. Proteins were extracted by adding 2 volumes of the following extraction buffer to 1 volume of grounded tissue: (50 mM Tris-HCl, pH 7.5, 150 mM NaCl, 10 % glycerol, 2.5 mM NaF, 2 mM NaMo, 1.5 mM activated Na3VO4, 5 mM DTT, 1% IGEPAL CA-630, 1x protease inhibitor cocktail 1 (Sigma Aldrich), and 1 mM PMSF) for 30 min at 4 °C with rotation. Samples were centrifuged at 4°C 4200g for 20 min and plant extract was passed through 4x Miracloth. Inputs were taken and kept on ice. 40 μM of anti-FLAG (or anti-GFP) protein beads equilibrated in extraction buffer were added to the rest of the extract. Protein immuno-precipitation was carried out for 2h at 4 °C with rotation. The beads were collected by centrifugation at 200 g for 2 min at 4 °C and washed 3 times in extraction buffer and twice in extraction buffer with 0.5 % IGEPAL CA630. Fifty microliters of 2x SDS buffer was added to each sample; the corresponding amount of 4x SDS buffer was added to input samples and all samples were heated at 70 °C for 10 min. Samples were either straight loaded on the gel or kept at −20.

### LC-MS/MS analysis

*35S::PUB4-FLAG* or Col-0 plants were grown the same way as for CoIP procedure. Ten milliliters of grounded plant tissue was used for one IP. The IP procedure was the same as described above for CoIP. Samples were run on SDS-polyacrylamide gel, and gel was stained with Coomassie Brilliant Blue (Simply BlueTM Safe stain, Invitrogen). Afterwards, each gel line was cut in several pieces, and these pieces were kept in separate 2 mL low protein binding tubes (Eppendorf). Afterwards, the samples were processed as described previously (Bender et al., 2017). Briefly, gel slices were de-stained in 50 % acetonitrile and incubated for 45 min in 10 mM DTT. Cysteinyl residue alkylation was carried out for 30 min in the darkness in 55 mM chloroacetamide. After several washes with 25 mM ammonium bicarbonate, 50 % acetonitrile gel slices were dehydrated in 100% acetonitrile. Gel pieces were rehydrated with 50 mM ammonium bicarbonate and 5% acetonitrile containing 20 ng/μL trypsin (Pierce), and digestion was performed overnight at 37 °C. Tryptic peptides were sonicated from the gel in 5 % formic acid, 50 % acetonitrile, and the total extracts were evaporated until dry.

LC-MS/MS analysis was performed using an Orbitrap Fusion trihybrid mass spectrometer (Thermo Fisher Scientific) and a nanoflow-UHPLC system (Dionex Ultimate3000, Thermo Fisher Scientific). Peptides were trapped to a reverse phase trap column (Acclaim PepMap C18, 5 μm, 100 μm × 2 cm, Thermo Fisher Scientific) connected to an analytical column (Acclaim PepMan 100, C18 3 μm, 75 μm × 50 cm, Thermo Fisher Scientific). Peptides were eluted in a gradient of 3-30 % acetonitrile in 0.1 % formic acid (solvent B) over 50 min followed by gradient of 30-80 % B over 6 min at a flow rate of 300 nL/min at 40 °C. The mass spectrometer was operated in positive ion mode with nano-electrospray ion source with an inner diameter of 0.02 mm fused silica emitter (New Objective). Voltage 2200 V was applied via platinum wire held in PEEK T-shaped coupling union with transfer capillary temperature set to 275 °C. The Orbitrap, MS scan resolution of 120,000 at 400 m/z, range 300 to 1800 m/z was used, and automatic gain control was set to 2 × 10^5^ and maximum inject time to 50 ms. In the linear ion trap, product ion spectra were triggered with a data-dependent acquisition method using “top speed” and “most intense ion” settings. The threshold for collision-induced dissociation (CID) and high energy collisional dissociation (HCD) was set using the Universal Method (above 100 counts, rapid scan rate, and maximum inject time to 10 ms). The selected precursor ions were fragmented sequentially in both the ion trap using CID and in the HCD cell. Dynamic exclusion was set to 30 s. Charge state allowed between +2 and +7 charge states to be selected for MS/MS fragmentation.

Mascot generic files (.mgf files) were generated from raw data using MSConvert package (Matrix Science) and were searched on Mascot server version 2.4.1 (Matrix Science) against TAIR (version 10) database, a separate in-house constructs database and an in-house contaminants database. Tryptic peptides with up to two possible mis-cleavages and charge states +2, +3, +4 were allowed in the search. The following modifications were included in the search: oxidized Met, phosphorylation on Ser, Thr, Tyr as variable modifications, and carbamidomethylated Cys as a static modification. Data were searched with a monoisotopic precursor and fragment ions mass tolerance 10 ppm and 0.6 Da, respectively. Mascot results were combined in Scaffold version 4 (Proteome Software) and exported to Excel (Microsoft Office).

### Parallel Reaction Monitoring analyses

Parallel reaction monitoring was performed as described in Guo et al., (2020). Briefly, phospho-peptides were targeted to measure PUB4 phosphorylation at indicated residues (Supplementary Table S3). The PRM assay also included a selection of non-modified control peptides (Supplementary Table S3) to measure PUB4 protein levels. These were used to normalize the measured changes in phosphorylation relative to PUB4 protein levels. For some phospho-peptides transitions did not identify the specific site of phosphorylation and could only be narrowed down regions (Figure 8 (B-E) and Supplementary Table S3). The assay was performed three times for each of two biological replicates and results averaged ± SE.

### Yeast two-hybrid screen

Yeast two-hybrid screening was conducted by Hybrigenics Services, S.A.S., Paris, France (http://www.hybrigenics-services.com). The RipAC coding region from *R. solanacearum* GMI1000 was PCR-amplified and inserted into pB29 as a N-terminal fusion to LexA DNA-binding domain (RipAC-LexA). The construct was confirmed by sequencing the full-length RipAC and used as a bait to screen a random-primed Tomato Infected Roots cDNA library constructed into pP6. pB29 and pP6 derive from the original pBTM116 (Vojtek and Hollenberg, 1995) and pGADGH (Bartel et al., 1993) plasmids, respectively. 133 million clones (more than 10-fold the complexity of the library) were screened using a mating approach with YHGX13 (Y187 ade2-101::loxP-kanMX-loxP, matα) and L40ΔGal4 (matα) yeast strains as previously described (Fromont-Racine et al., 1997). Twenty-six His+ colonies were selected on a medium lacking tryptophan, leucine and histidine. The prey fragments of the positive clones were amplified by PCR and sequenced at their 5’ and 3’ junctions. The resulting sequences were subjected to corresponding interacting proteins analysis in the GenBank database (NCBI) using a fully automated procedure.

### Split-LUC assays

Split-LUC assays were performed as previously described (Chen et al., 2008; Yu et al., 2020). Generally, *A. tumefaciens* strains containing the desired plasmids were infiltrated into *N. benthamiana* leaves. Split-LUC assays were conducted both qualitatively and quantitatively after 2 dpi. For the CCD imaging, the leaves were infiltrated with 0.5 mM luciferin in water and kept in the dark for 5 min before CCD imaging. The images were taken with either Lumazone 1300B (Scientific Instrument, West Palm Beach, FL, US). To perform the quantification of the luciferase signal, leaf discs (diameter=4 mm) were taken into a 96-well microplate (PerkinElmer, Waltham, MA, US) with 100 ◻L H_2_O. Then the leaf discs were incubated with 100 ◻L water with 0.5 mM luciferin in a 96-well plate wrapped with foil paper to remove the background luminescence for 5 min, and the luminescence was recorded with a Microplate luminescence reader (Varioskan flash, Thermo Scientific, USA). Each data point contains at least eight replicates. The protein accumulation was determined by immunoblot as described above.

### FRET-FLIM

F◻rster resonance energy transfer – fluorescence lifetime imaging (FRET-FLIM) experiments were performed as previously described (Rosas-Diaz et al., 2018; Xian et al., 2019) with several modifications. Briefly, AtPUB4 and SlPUB4 (fused to GFP) were expressed from epiGreen-35S and pGWB505, respectively; and RipAC (fused to RFP) was expressed from pGWB554. FRET-FLIM experiments were performed on a Leica TCS SMD FLCS confocal microscope excitation with WLL (white light laser) and emission collected by a SMD SPAD (single photon-sensitive avalanche photodiodes) detector. Two days after infiltration, *N. benthamiana* plants transiently coexpressing donor and acceptor proteins were visualized under the microscope. Accumulation of the GFP- and RFP-tagged proteins was estimated before measuring lifetime. The tuneable WLL set at 488 nm with a pulsed frequency of 40 MHz was used for excitation, and emission was detected using SMD GFP/RFP Filter Cube (with GFP: 500-550 nm). The fluorescence lifetime shown in the figures corresponding to the average fluorescence lifetime of the donor was collected and analyzed by PicoQuant SymphoTime software. Lifetime is normally amplitude-weighted mean value using the data from the single (GFP-fused donor protein only or GFP-fused donor protein with free RFP acceptor or with non-interacting RFP-fused acceptor protein) or biexponential fit (GFP-fused donor protein interacting with RFP-fused acceptor protein). Mean lifetimes are presented as mean ± SEM based on eight images from three independent experiments.

### Tomato root transformation

Tomato root transformation was performed as previously described (Morcillo et al., 2020). Briefly, the radicle and bottom part of the hypocotyl of four-days-old tomato seedlings were removed and cut seedlings were immersed on *Agrobacterium rhizogenes* MSU440 bacterial mass containing pUBIcGFP-DR::SlPUB4 for overexpression and pK7GWIWG2_II-RedRoot::SlPUB4 for RNAi silencing (pUBIcGFP-DR and pK7GWIWG2_II-RedRoot empty vector were used as control). Seedlings inoculated with *A. rhizogenes* were kept on half-strength MS medium (0.8 % agar) and covered with filter paper to maintain humidity. During the following weeks, roots were screened and selected by DsRed fluorescence visualization, using In Vivo Plant Imaging System NightShade LB 985 (Berthold Technologies). Subsequent plant handling and *R. solanacearum* inoculation was performed as previously described (Morcillo et al., 2020).

### Quantification and statistical analysis

Statistical analyses were performed with Prism 7 software (GraphPad). The data are presented as means ± SE. The statistical analysis methods are described in the figure legends.

## Supplementary Tables

**Suppementary Table S1.**
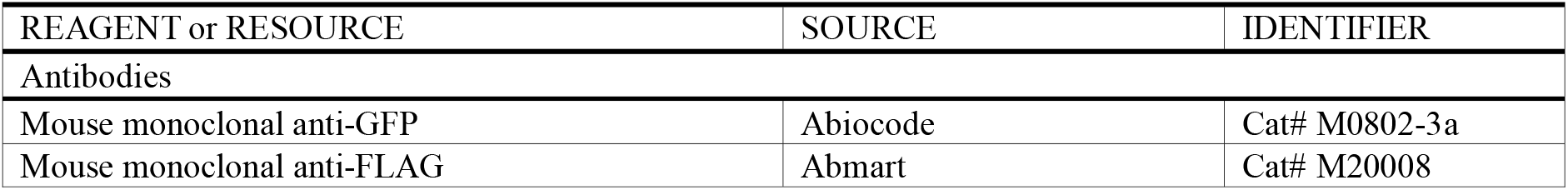
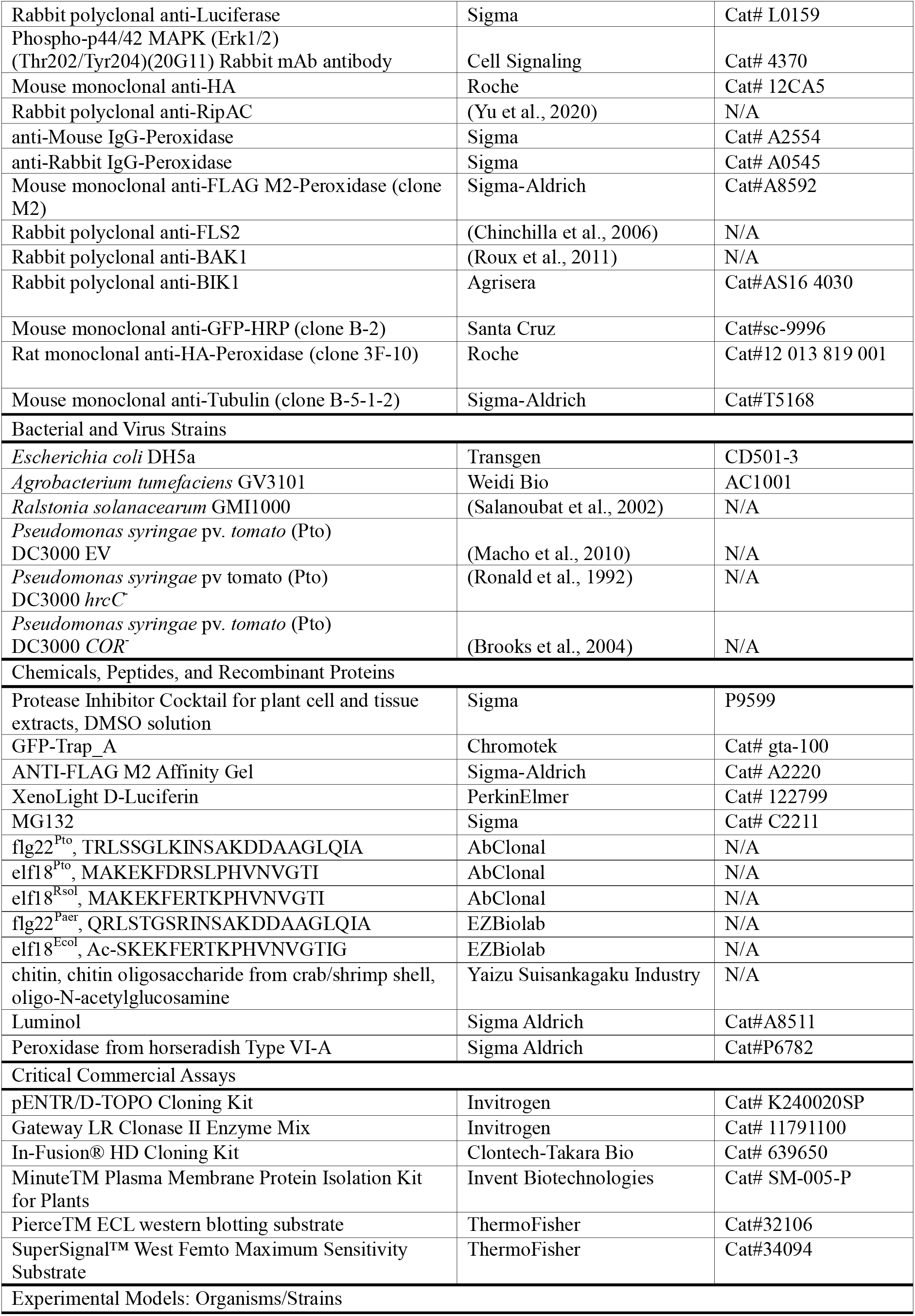
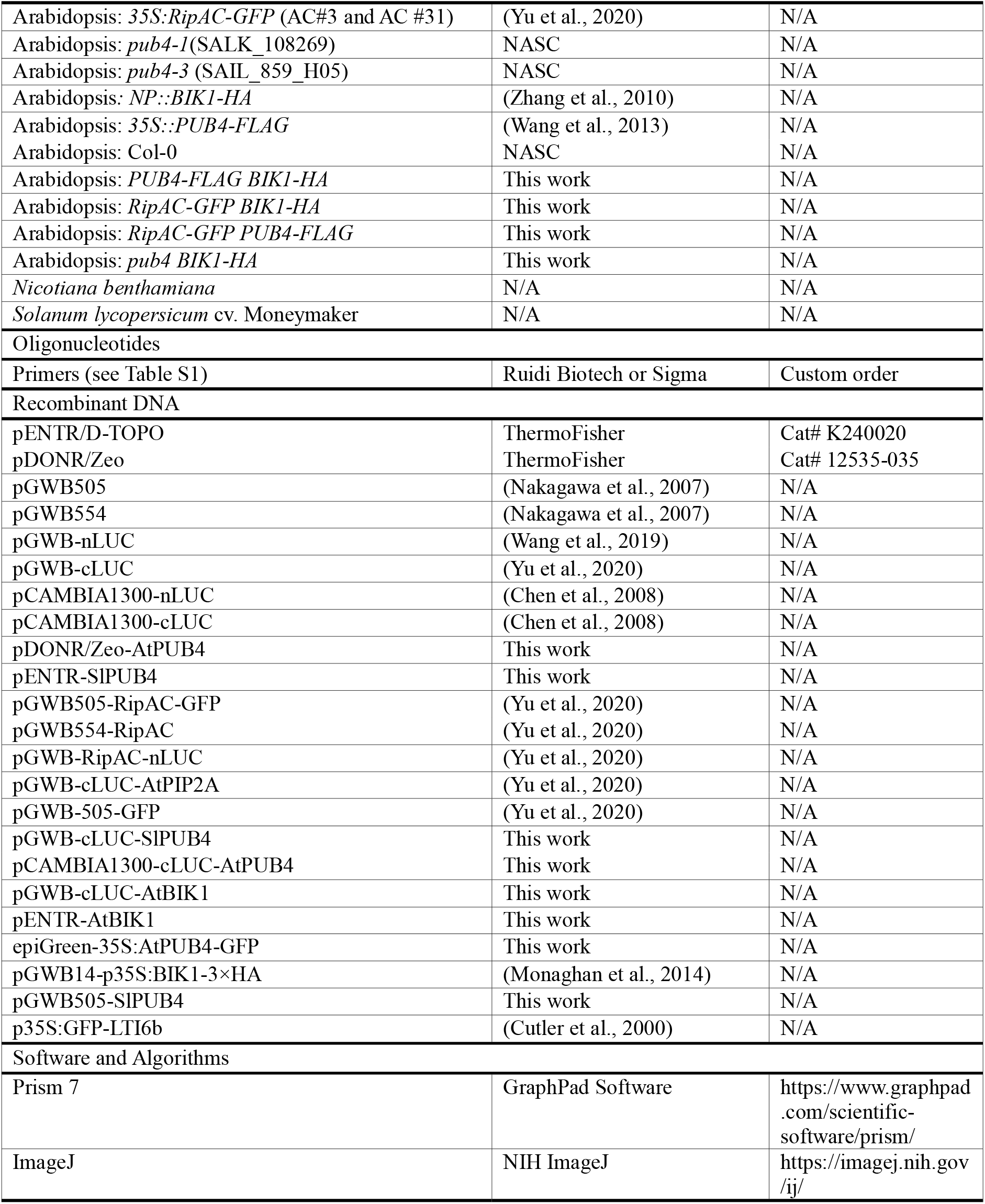
Resources used in this study.

**Suppementary Table S2.**
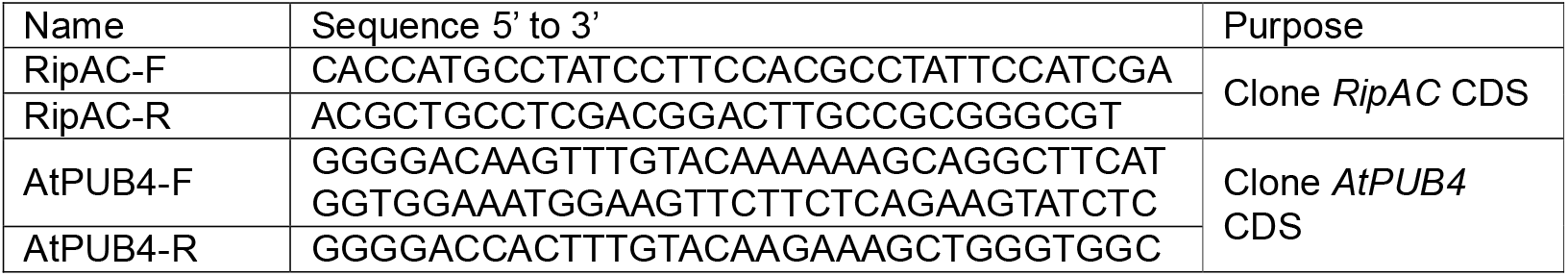
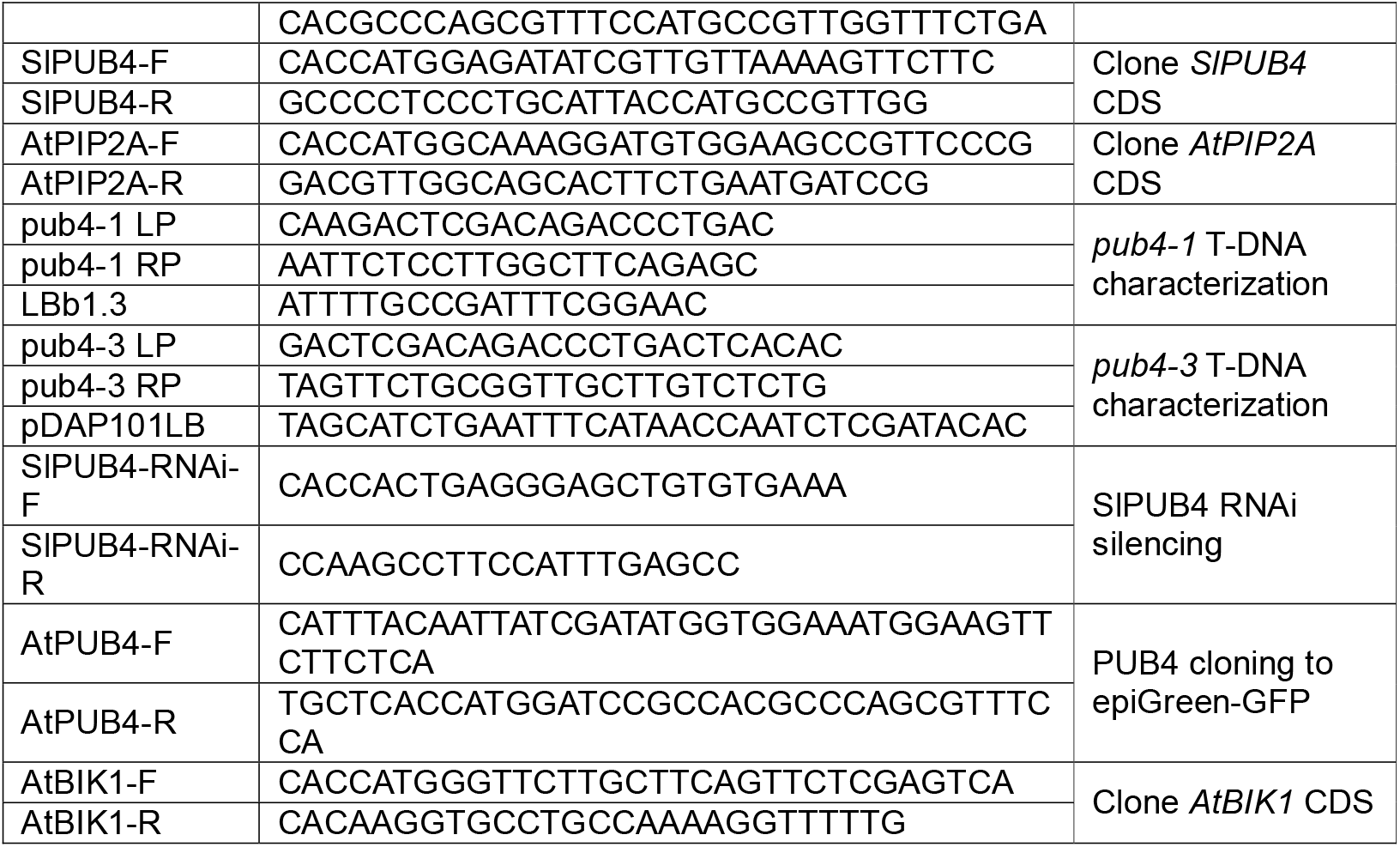
Primers used in this study.

**Suppementary Table S3.**
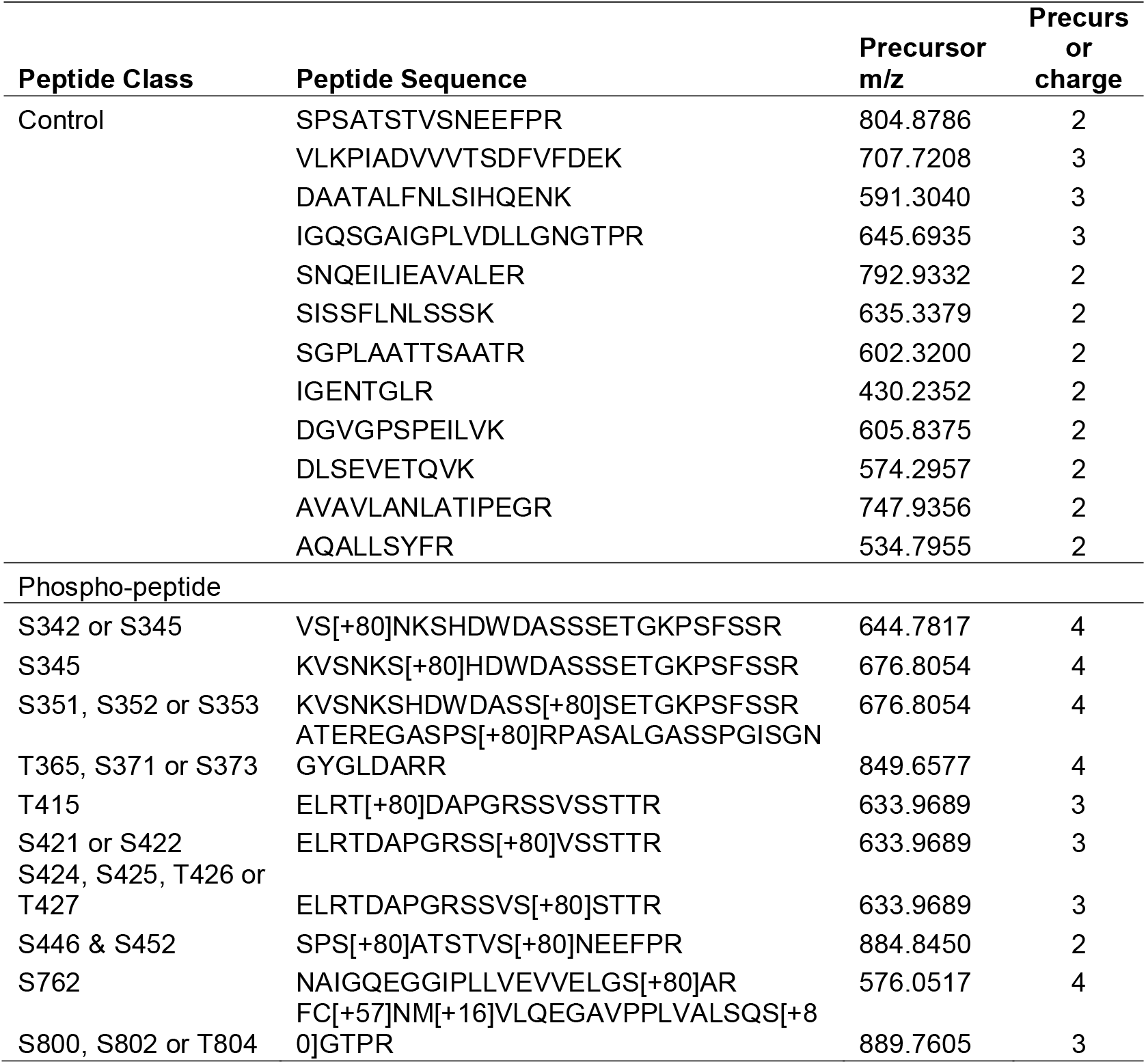
Peptide m/z and charge state used to target PUB4 control and phospho-peptides.

**Figure S1.**
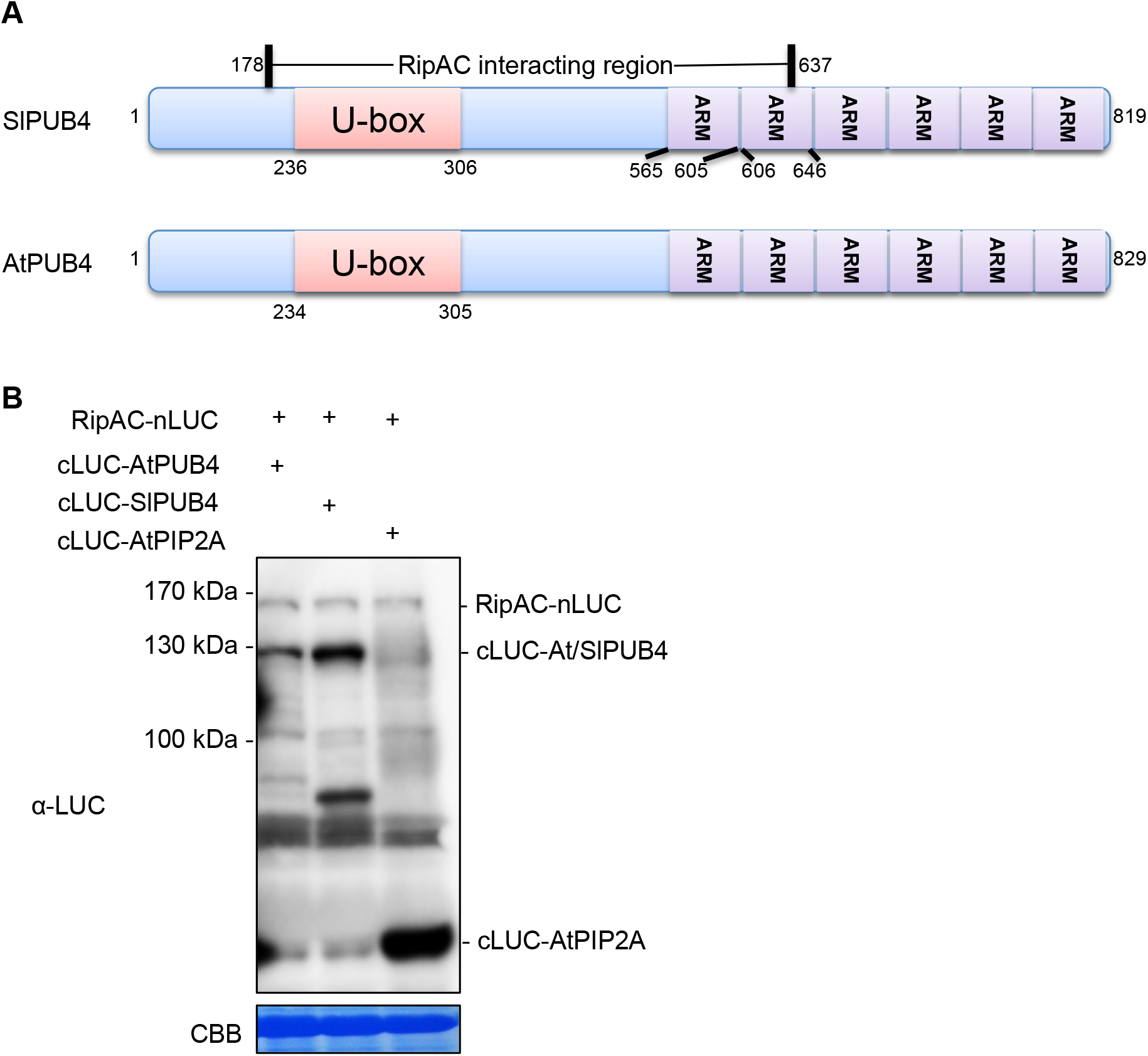
RipAC associates with PUB4 in plant cells. (A) Yeast two-hybrid screening identified SlPUB4 as RipAC-interacting protein. The diagrams depict domain architectures of SlPUB4 and AtPUB4, while RipAC interacting region in SlPUB4 through yeast two-hybrid is labeled. ARM means Armadillo domain. (B) Western blot showing protein accumulation in the experiments shown in Figures 2A and 2B. Immunoblots were analyzed using anti-luciferase (LUC) antibody. Coomassie brilliant blue (CBB) staining was used as loading control. Molecular weight (kDa) marker bands are indicated for reference.

**Figure S2.**
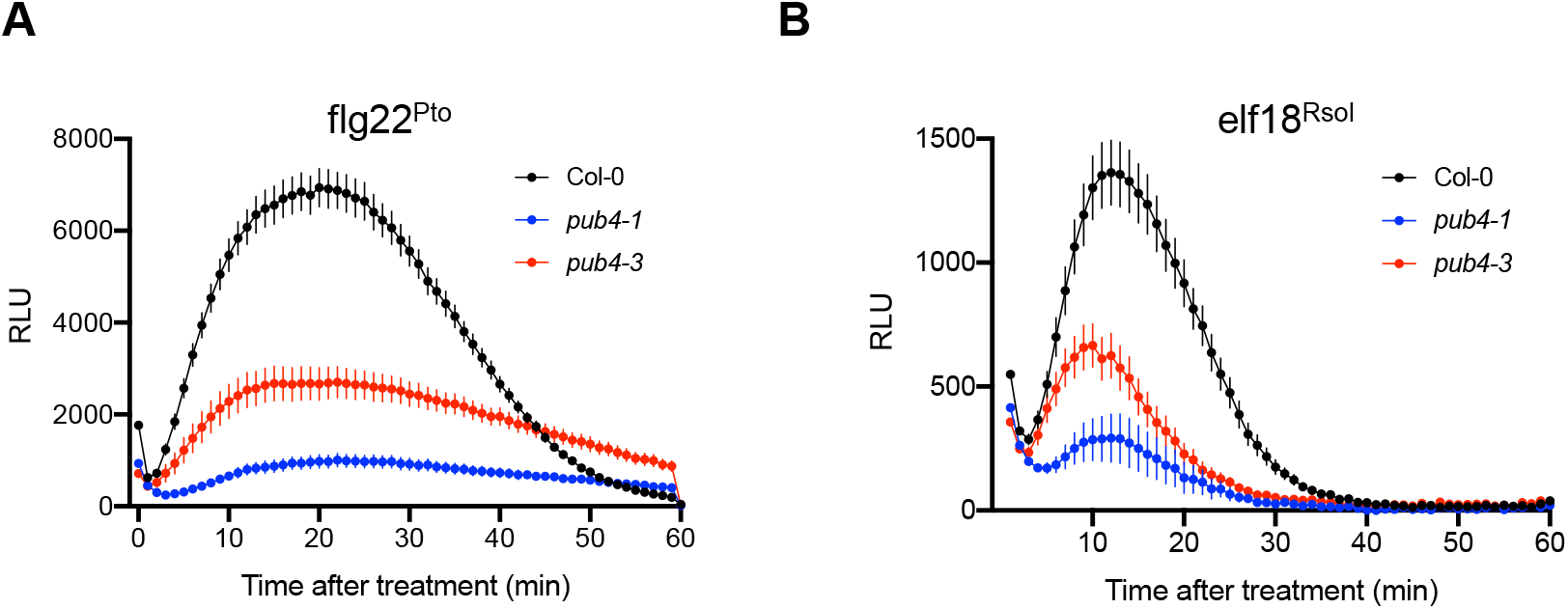
PUB4 positively regulates PTI in Arabidopsis. ROS burst in Col-0 WT or the indicated *pub4* mutant lines induced by 100 nM flg22 from *Pto* DC3000 (flg22^Pto^) (A) or 100 nM elf18 from *R. solanacearum* (elf18^Rsol^) (B). ROS was measured as relative luminescence units (RLU) over time. Values are means ± SE (n=16). Experiments were performed at least three times with similar results.

**Figure S3.**
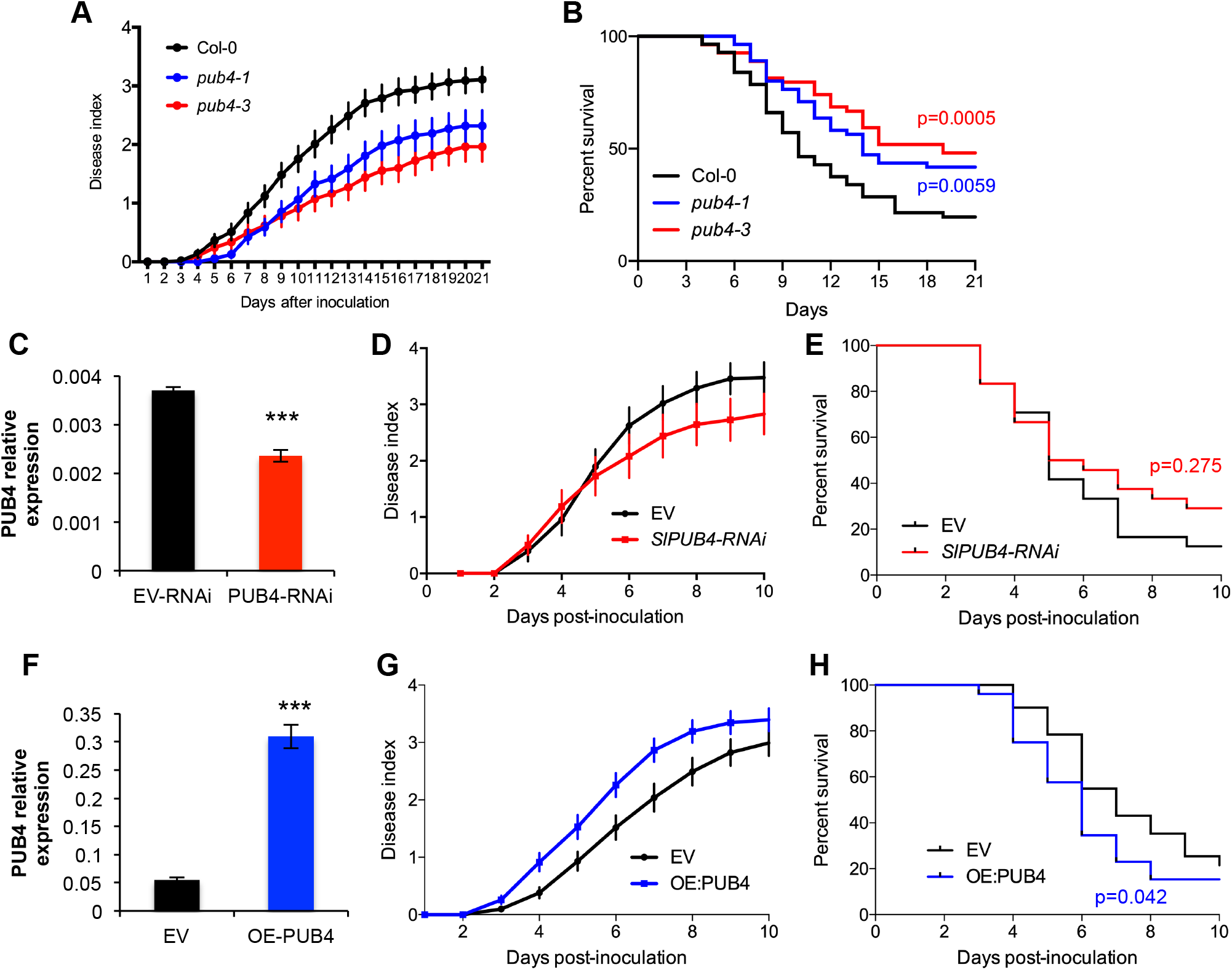
PUB4 promotes *R. solanacearum* infection in Arabidopsis and tomato. (A and B) Arabidopsis *pub4* mutants show enhanced resistance to *R. solanacearum* infection. Composite data from 3 independent biological repeats (a representative assay is shown in Figure 4A). All values were pooled together and represented as disease index (A) or percent survival (B). Disease index values represent means ± SE (n=55). To calculate the percentage of survival, the disease scoring was transformed into binary data with the following criteria: a disease index lower than 2 was defined as ‘0’, while a disease index equal or higher than 2 was defined as ‘1’ for each specific time point. Statistical analysis was performed using a Log-rank (Mantel-Cox) test, and the corresponding p value is shown in the graph with the same colour as each curve. (C) Expression of the *SlPUB4* gene in tomato roots expressing the *SlPUB4*-RNAi construct used in the experiments shown in Figure 4B, determined by qRT-PCR. Values were normalized to the expression of the *SlEF1α-1* gene, and are shown as relative to the expression in roots expressing the empty vector (EV). Values represent means ± SE (n=3 samples per genotype), and asterisks represent significant differences according to a Student’s t test (***p<0.001). (D and E) Tomato plants expressing the *SlPUB4*-RNAi construct show enhanced resistance to *R. solanacearum* infection. Composite data from 3 independent biological repeats (a representative assay is shown in Figure 4B). All values were pooled together and represented as disease index (D) or percent survival (E). Disease index values represent means ± SE (n=24). To calculate the percentage of survival, the disease scoring was transformed into binary data with the following criteria: a disease index lower than 2 was defined as ‘0’, while a disease index equal or higher than 2 was defined as ‘1’ for each specific time point. Statistical analysis was performed using a Log-rank (Mantel-Cox) test, and the corresponding p value is shown in the graph with the same colour as each curve. (F) Expression of the *SlPUB4* gene in tomato roots overexpressing *SlPUB4* used in the experiments shown in Figure 4C, determined by qRT-PCR. Values were normalized to the expression of the *SlEF1α-1* gene, and are shown as relative to the expression in roots expressing the empty vector (EV). Values represent means ± SE (n=3 samples per genotype), and asterisks represent significant differences according to a Student’s t test (***p<0.001). (G and H) Tomato plants with roots overexpressing *SlPUB4* show enhanced susceptibility to *R. solanacearum* infection. Composite data from 7 independent biological repeats (a representative assay is shown in Figure 4C). All values were pooled together and represented as disease index (G) or percent survival (H). Disease index values represent means ± SE (n=52). To calculate the percentage of survival, the disease scoring was transformed into binary data with the following criteria: a disease index lower than 2 was defined as ‘0’, while a disease index equal or higher than 2 was defined as ‘1’ for each specific time point. Statistical analysis was performed using a Log-rank (Mantel-Cox) test, and the corresponding p value is shown in the graph with the same colour as each curve.

**Figure S4.**
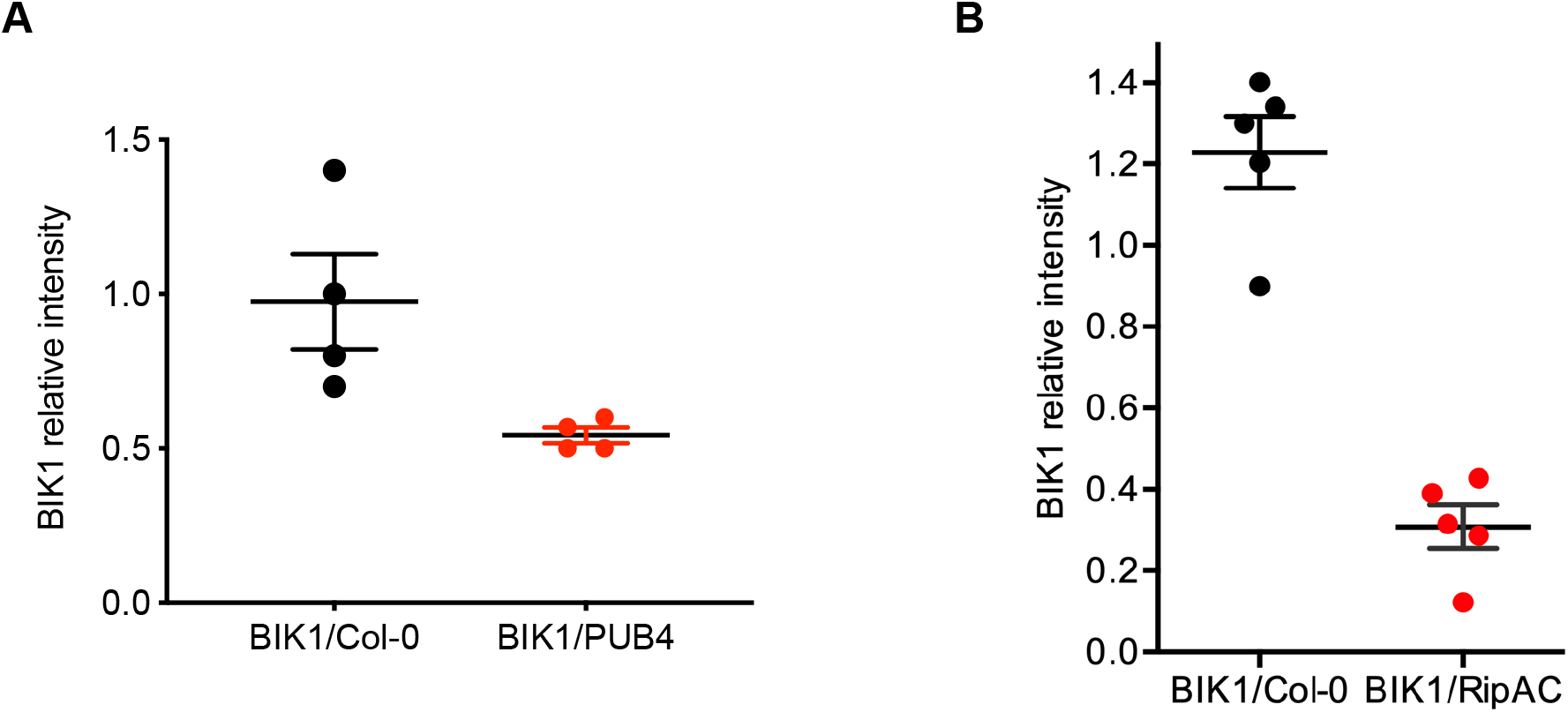
PUB4 and RipAC reduce BIK1 protein accumulation. (A) Quantification of relative protein abundance from the western blot in Figure 7C, using ImageJ software. Relative intensity of BIK1-HA compared with tubulin is represented for each sample and corresponding genotype. (B) Quantification of relative protein abundance from the western blot in Figure 7F. Protein band intensities were quantified using ImageJ software. Relative intensity of BIK1-HA compared with actin is represented for each sample and corresponding genotype. Experiments were performed at least three times with similar results.

**Figure S5.**
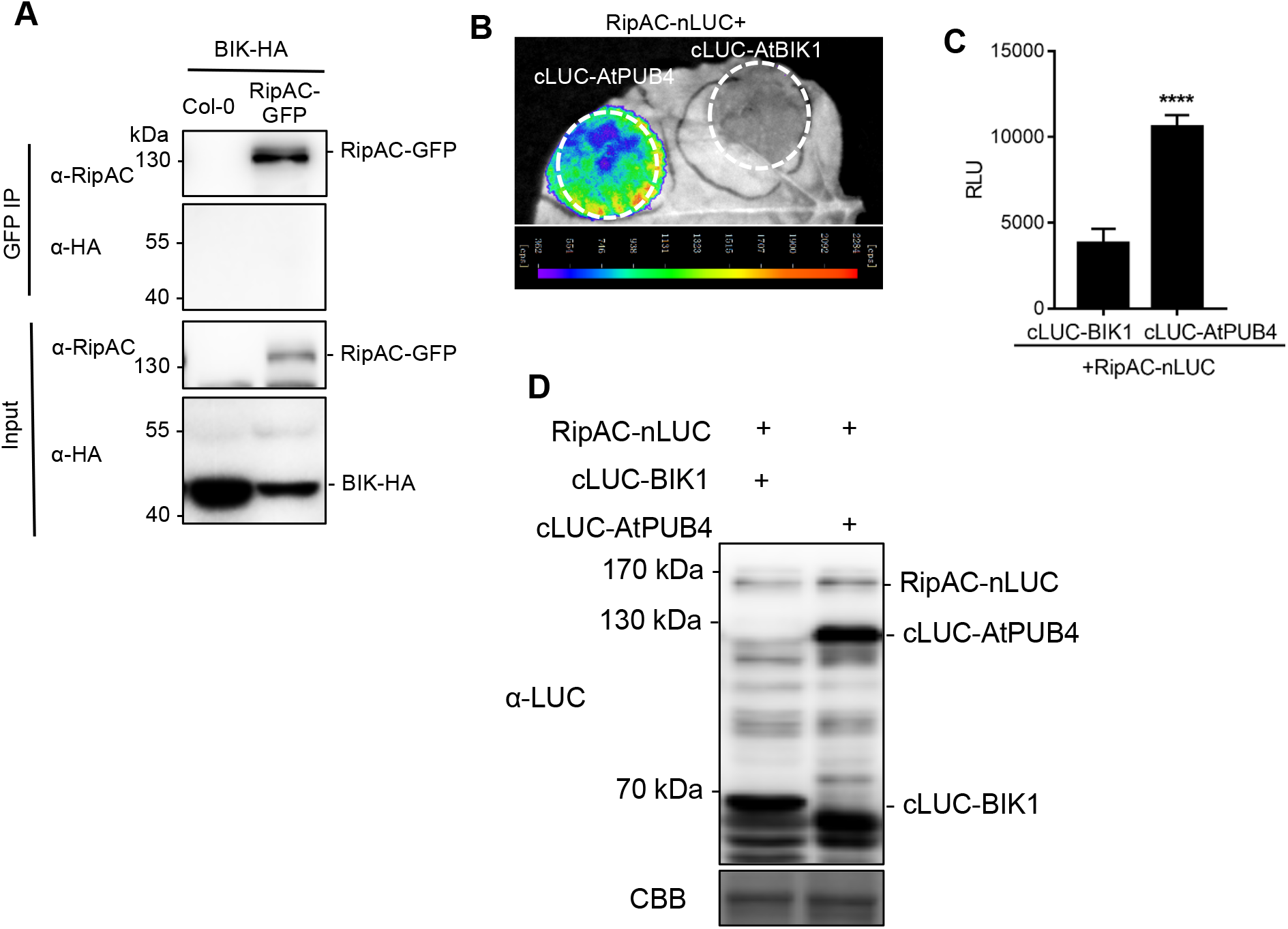
RipAC does not associate with BIK1 *in planta*. (A) Co-immunoprecipitation of RipAC-GFP and BIK1-HA was performed in RipAC-GFP/BIK1-HA F2 progeny pool, while Col-0 crossed with BIK1-HA was used as a control. Twelve days after germination seedling tissues were collected and then subjected to anti-GFP immunoprecipitations. In IP, anti-HA blots were developed using Femto substrate. Three independent biological replicates were performed. (B-D) RipAC does not interact with BIK1 in Split-LUC assays, either qualitatively (B) or quantitatively (C), in *Nicotiana benthamiana*. The interaction with PUB4 (also shown in Figure 2) was used as positive control. Values are means ± SE (n=8). Asterisks indicate significant differences between both samples (Student’s t test, **** p<0.0001). (D) Western blot showing protein accumulation in these tissues. Immunoblots were analyzed using anti-luciferase (LUC) antibody. Coomassie brilliant blue (CBB) staining was used as loading control. Molecular weight (kDa) marker bands are indicated for reference.

**Figure S6.**
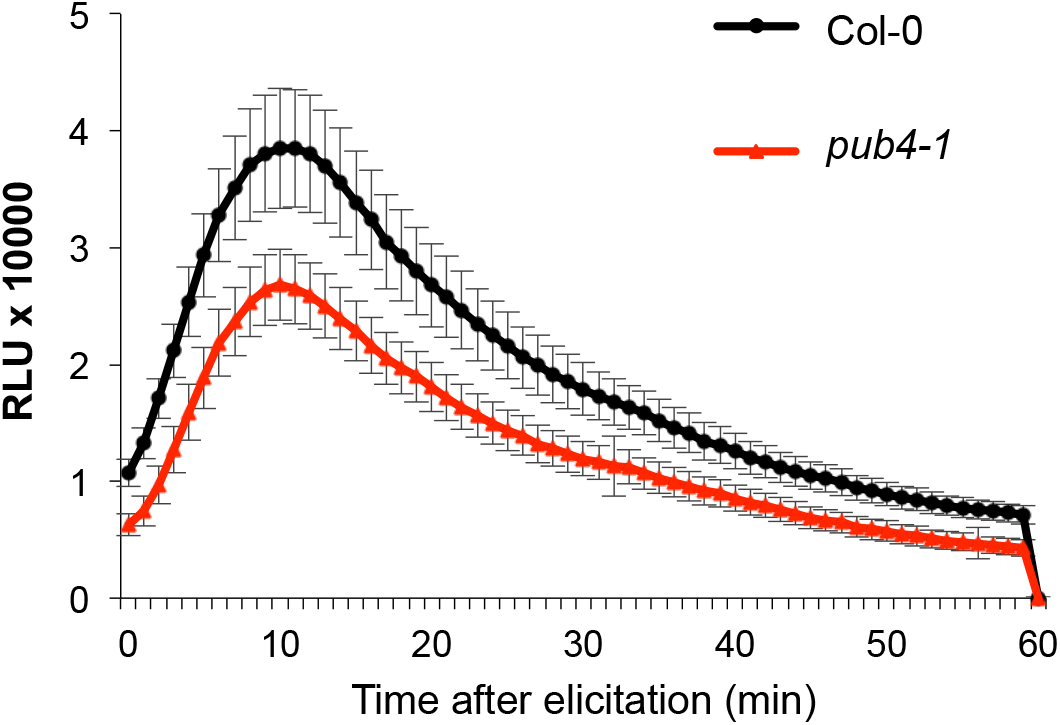
The *pub4-1* mutant shows reduced flg22-triggered ROS in Arabidopsis roots. ROS production in roots of Col-0 WT or the *pub4-1* mutant, induced by 100 nM flg22. ROS was measured as relative luminescence units (RLU) over time. Values are means ± SE (n=16). Experiments were performed three times with similar results.

